# Regulation of *KRAS4A/B* splicing in cancer stem cells by the RBM39 splicing complex

**DOI:** 10.1101/646125

**Authors:** Wei-Ching Chen, Minh D. To, Peter M.K. Westcott, Reyno Delrosario, Il-Jin Kim, Mark Philips, Quan Tran, Nora Bayani, Allan Balmain

**Author notes:** These authors contributed equally to this work (Wei-Ching Chen, Minh D. To, Peter M.K. Westcott).

## Abstract

The *KRAS* oncogene is expressed as major KRAS4B and minor KRAS4A splice isoforms, the distinct functions of which in cancer are unknown. We demonstrate here that KRAS4A is enriched in cancer stem-like cells, and is activated by hypoxia, whereas KRAS4B is more widely expressed and responds to ER stress. Mice completely lacking either isoform are viable but resistant to lung cancer development, as are *Kras4A/B* double heterozygous mice expressing both isoforms, but in which splice regulation has been uncoupled. Splicing of KRAS4A, but not KRAS4B, in human tumor cells can be inhibited by treatment with the splice inhibitor indisulam, or by CRISPR/Cas inhibition of the RBM39 splicing complex. Our data suggest that control of *KRAS4A/B* splicing is a targetable vulnerability in *KRAS* mutant tumors.

---

*KRAS* undergoes alternative splicing of the last exon to generate two proteins, KRAS4A and KRAS4B, which are identical except for the 22/23 amino acids at the carboxyl terminus required for post-translational modifications and intracellular trafficking(*1*). Studies in the mouse have shown that germline deletion of the complete *Kras* gene results in embryonic lethality (*2*), but specific deletion of only *Kras4A*, achieved by deletion of exon 4A(*3*) or knock-in of *Kras4B* cDNA (*Kras4B^KI^* mice, henceforth *Kras4A^−/−^*) (*4*), had no effect on viability, suggesting that the main developmental functions of *Kras* are mediated through the *Kras4B* isoform. It has previously been demonstrated that mice lacking only the *Kras4A* isoform are resistant to chemically induced lung cancer(*5, 6*), in spite of the fact that *Kras4A* is expressed only in a subpopulation of normal and tumor cells. These data led to the proposal that Kras4A plays an essential role in tumor development, possibly through effects on a minor stem cell population(*5*). Subsequent analysis of human cancer data from TCGA also suggested a more important role for KRAS4A in human cancer than had previously been appreciated, as some tumors expressed unexpectedly high levels of this “minor” isoform(*7*).

We designed a genetic approach to investigate the distinct functions of Kras4A and Kras4B in mouse models and human cancers. We show that each isoform is individually dispensable for normal mouse development, and that coordinated expression of both isoforms is required for initiation and growth of *Kras* mutant cancers. Inactivation of *Kras4A* in mouse and human cells reveals distinct functions of *Kras4A* in stress responses and tumor metabolism, particularly in a sub-population of cells with stem cell properties. Our data suggest that coordination of these different functions in cancers may be achieved through regulation of mRNA splicing, identifying a potential novel route to targeting the *KRAS* pathway during tumor development.

## Results

### The major Kras4B isoform is dispensable for normal mouse development

To investigate the specific role of *Kras4B* in development and lung carcinogenesis *in vivo*,we used the previously published strategy(*4*) to insert a *Kras4A* cDNA into the *Kras* locus, thus generating an allele that completely lacks expression of *Kras4B* (*Kras4B^−/−^* mice). Specifically, *Kras4A* cDNA spanning a portion of exon 2 through 3’ UTR was homologously inserted in-frame between exons 2 and 3 of the *Kras* locus (Fig. S1A). PCR and Southern blot analysis confirmed the correct homologous targeting of this construct (Fig. S1B-C).

*Kras4B^−/−^* homozygous mice were born at normal Mendelian ratios, although they showed a slight but significant reduction in weight at weaning compared to WT littermates (Fig. S1D). In contrast with previously generated *Kras4A^−/−^* mice, Kras4B^−/−^ animals were infertile. Normal skin, lung, and heart of WT animals showed minimal Kras4A and high Kras4B protein levels, while normal colon and kidney showed high expression of both, consistent with reported expression patterns in these tissues(*8*) (Fig. S1E). As anticipated, these tissues showed intermediate levels of Kras4A and Kras4B proteins in *Kras4B^+/−^* heterozygous animals, and only Kras4A in homozygous animals. Furthermore, Kras4A showed incremental increases in *Kras4B^+/−^* heterozygous *and Kras4B^−/−^* homozygous mice, with levels in the skin, lung, and heart—normally very low—markedly higher in *Kras4B^−/−^* animals for all tissues analyzed, within expected inter-animal variation (Fig. S1E).

To investigate Ras signaling through the canonical Mapk and Akt pathways, primary mouse embryonic fibroblasts (MEFs) isolated from E13.5 embryos were serum starved overnight prior to stimulation with EGF for different lengths of time. We observed a robust EGF-induced activation of MapK and Akt that peaked at the 5-minute time point in all three genotypes, suggesting that lack of Kras4A or Kras4B did not affect the response of these canonical RAS signaling pathways to EGF stimulation. Moreover, using the Raf RBD assay to capture active GTP-bound Ras proteins, we demonstrated that Kras4A and Kras4B are both activated in response to EGF in cells that lack the alternative isoform (Fig. S2).

### *Both Kras4A* and *Kras4B* are essential for mouse lung tumorigenesis

In order to compare the roles of *Kras4A* and *Kras4B* in lung tumorigenesis, both knockout strains were crossed over 6 generations onto the *FVB/N* background, and injected with urethane. Contrary to our expectations, we found that *Kras4B^−/−^* homozygous mice were highly resistant to lung tumor development (Fig. 1A-D). Of the 16 *Kras4B^−/−^* homozygous animals treated with urethane, 8 developed one or two extremely small tumors each (Fig. 1A-B), and Sanger sequencing of *Kras* codons 12, 13, and 61 revealed no mutations. The lack of tumors in *Kras4A^−/−^* animals agrees with previously published results(*5, 6*). Thus, the effect of eliminating *Kras4B*, similar to loss of *Kras4A*, is to confer resistance to chemically-induced lung tumours with *Kras* mutations.

**Figure 1.**
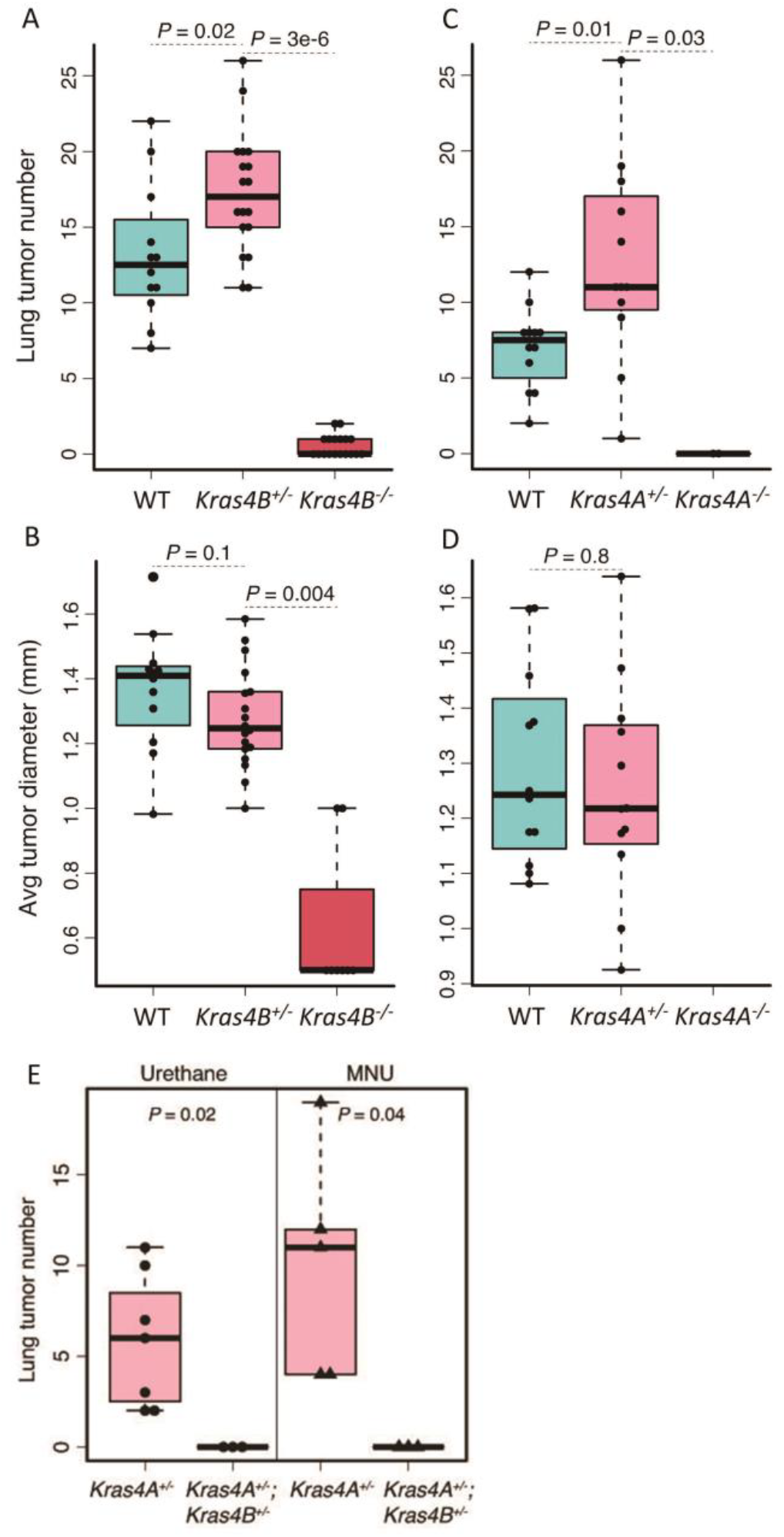
Kras4A and Kras4B are essential for development of carcinogen-induced lung tumours. Urethane-induced lung tumor number (A) and average diameter (B) in *WT*, *Kras4B^−^* heterozygous and homozygous mice. Homozygous mice showed a highly significant reduction in tumor number and size, while heterozygous mice developed significantly more tumors than *WT* mice. (C-D) Urethane-induced lung tumor number (C) and average diameter (D) in *WT*, *Kras4A^−^* heterozygous and homozygous mice showed very similar patterns as seen in the *Kras4B^−^* cross. (E) Urethane-induced (left panel) and MMU-induced (right panel) lung tumor number in *Kras4A^+/−^* heterozygous and double heterozygous *Kras4A^+/−^, Kras4B^+/−^* mice. No tumors were found in double heterozygous *Kras4A^+/−^, Kras4B^+/−^* mice. All lung tumorigenesis experiments were performed on a *FVB/N* background.

Any tumors that arose in either *Kras4A^+/−^* or *Kras4B^+/−^* heterozygous mice carried mutations in the fully functional wild type allele (Table S1), suggesting that the coordinated activity of mutant forms of both isoforms may be essential for Kras-driven tumorigenesis. To test the possibility that a mutation in one isoform could cooperate with a wild type alternative isoform, we carried out carcinogenesis studies in the lung using double heterozygous *Kras4A^+/−^,Kras4B^+/−^* mice. Specifically, 5 double heterozygotes were treated with 5 doses of urethane, 3 with 3 doses of urethane, and 5 with 3 doses of the carcinogen N-methyl-N-nitrosourea (MNU) in an attempt to increase the mutation burden and likelihood of generating mutant forms of one or both isoforms. No tumors were detected in the lungs of any of these double heterozygous animals 30 weeks after treatment (Fig. 1E). The resistance of mice expressing both isoforms, but on different alleles, suggests that the initiating *Kras* mutation has to be in an endogenous *Kras* gene capable of generating mutant versions of both splice variants. These data suggested that Kras4A and Kras4B splice isoforms may have complementary functions in ensuring tumor outgrowth from initiated cells. We therefore sought to identify functional differences between these isoforms that may contribute to their tumorigenic properties.

### Both *KRAS*4A and *KRAS4B* contribute to the tumorigenicity of *KRAS* mutant human cancers

To assess the functional relevance of the oncogenic KRAS isoforms in human cancer cell lines *in vitro*, we carried out shRNA-mediated knockdown and CRISPR/Cas knockout of each isoform in *KRAS* mutant cancer cell lines A549, SUIT2, YAPC and H358. We screened several hybridoma cell lines to identify antibodies capable of detecting endogenous KRAS4A (see Methods) which is normally expressed at low levels in cultured cells. Specific knock-down by isoform specific shRNA was confirmed by Westerns with the KRAS4A-specific antibody 10C11E4, and the KRAS4B-specific antibody (Santa Cruz Kras2b antibody) (Fig. S3A). Knockdown of either *KRAS4A* or *KRAS4B* in SUIT2 cells significantly compromised their capability to form colonies in soft-agar assays (Fig. S3B), suggesting that both KRAS4A and KRAS4B are required for growth.

We next used CRISPR/Cas technology to generate knockouts of *KRAS4A* or *KRAS4B* in human *KRAS* mutant cells. Single guide RNAs (sgRNAs) were designed to target the Cas9 nuclease to *KRAS* exons 4A or 4B in the human lung and pancreatic cancer cell lines A549 (G12S mutation) and SUIT2 (G12D mutation). Frameshift insertions and deletions in *KRAS* exon 4A or 4B were determined by allele specific cloning and sequencing of selected clones (Fig. S4). We were unable to isolate any homozygous *KRAS4B* frameshift mutant clones, and noted a greatly reduced number of colonies proliferating beyond the first few cell divisions in this arm of the experiment. Complete loss of KRAS4A and reduced levels of KRAS4B were shown using isoform-specific antibodies (Fig. 2A). To determine the effects of *KRAS* isoform-specific knockouts on cell growth, cell proliferation on plastic and in soft agar were assessed. Homozygous knockout of *KRAS4A* and heterozygous knockout of *KRAS4B* significantly reduced growth in both assays (Fig. 2B-D). Subcutaneously injected knockout cells in nude mice also had impaired tumor forming capacity compared to the parental SUIT2 or A549 cells (Fig. 2E). Taken together, these data indicate that KRAS4A and KRAS4B are both required for optimal growth of *KRAS* mutant lung and pancreatic tumor cells *in vitro* and *in vivo*.

**Figure 2.**
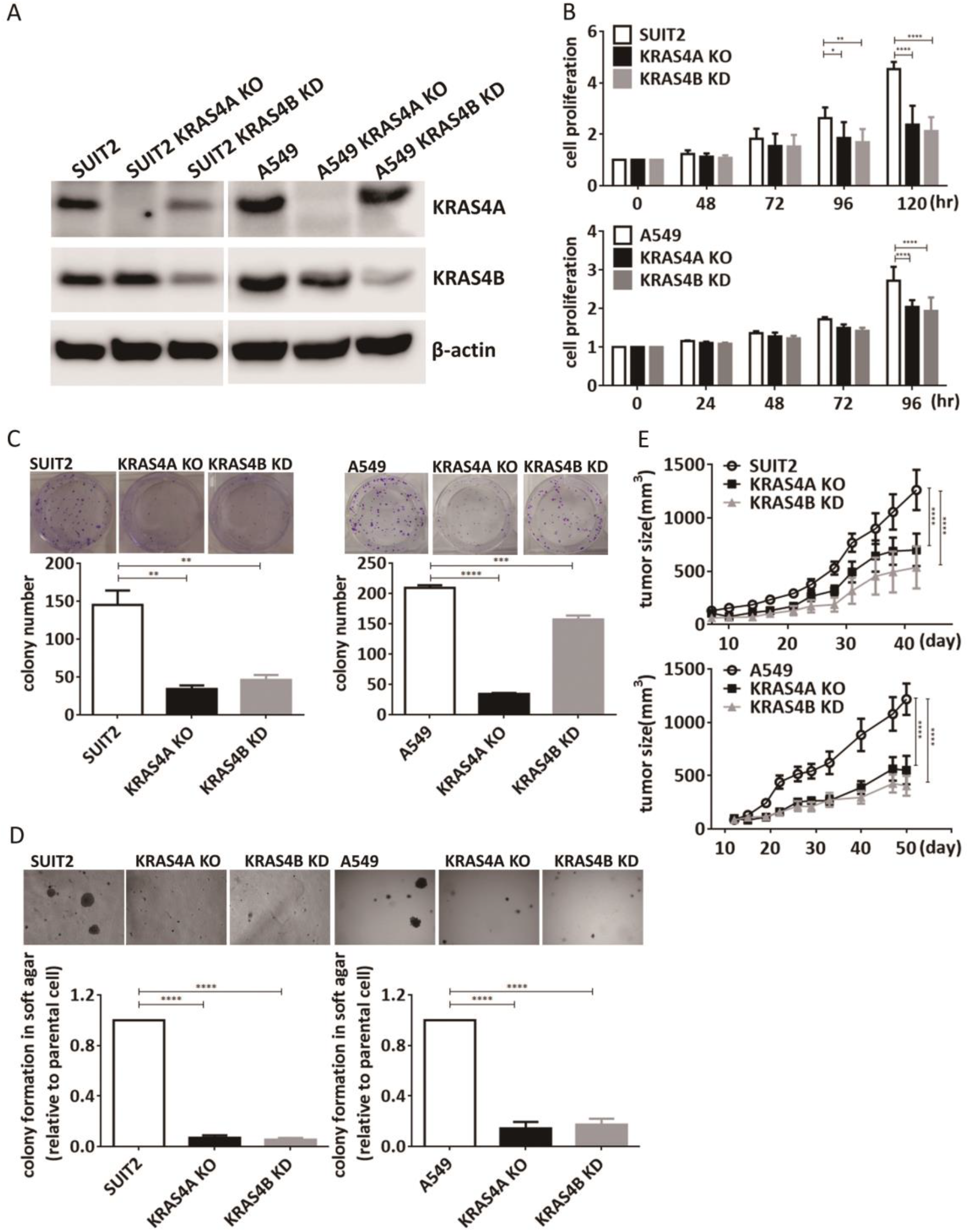
Effects of loss or reduced expression of KRAS4A or 4B on growth of human cancer cells *in vitro* and *in vivo*. (A) Western blotting using isoform-specific antibodies showed complete loss of KRAS4A (KRAS4A KO) in A549 and SUIT2 cells. KRAS4B is reduced but not eliminated in cells heterozygous for CRISPR/Cas-mediated heterozygous loss of *KRAS4B (KRAS4B KD)*. CRISPR-Cas9 induced knockouts of *KRAS4A* and *KRAS4B* in SUIT2 and A549 cells showed reduced growth *in vitro* on plastic (B), colony formation (C) and soft agar (D). Data are presented as mean ± s.e.m from three biological replicates. \**P* < 0.05; \*\**P* < 0.01; \*\*\**P* < 0.001; \*\*\*\**P* < 0.0001 by two-way ANOVA with Bonferroni’s multiple comparison for (B) and one-way ANOVA with Tukey’s multiple comparison for (C) and (D). (E) Growth of homozygous KRAS4A KO and heterozygous KRAS4B KD SUIT2 and A549 cells after subcutaneous injection into nude mice. Reduction in expression of both isoforms caused a significant decrease in tumor forming capacity *in vivo*. Data are presented as mean ± s.e.m. n = 10 mice for parental cell and n = 5 mice for KRAS4A KO or KRAS4B KD cell. \*\*\*\**P* < 0.0001 by two-way ANOVA.

We next investigated the relative expression levels of the two *KRAS* splice isoforms in *KRAS* mutant human lung cancer data sets (*9, 10*). We assessed the isoform-specific expression of *KRAS* in a cohort of 86 human lung adenocarcinomas from UCSF. Twenty three of these tumors harbor activating mutations in *KRAS* codons 12 or 13, while the remaining 63 are WT at *KRAS* codons 12, 13, and 61. Quantitative PCR analysis revealed that both total *KRAS* and *KRAS4A* expression are significantly elevated in the *KRAS* mutant subset of tumors, while *KRAS4B* expression showed no significant difference (Fig. S5A-C). We also analyzed the isoform-specific expression levels of *KRAS* from the TCGA lung adenocarcinoma (LUAD) RNA sequencing (RNAseq) dataset. Highly significant increases in both *KRAS4A* and *KRAS4B* transcripts were found in RNA sequencing (RNAseq) data from *KRAS* mutant versus WT lung adenocarcinoma (LUAD) tumors, but *KRAS4A* showed a greater fold increase (1.7) than *KRAS4B* (1.2) (Fig. S5D-E). To assess the alternative splicing of *KRAS* in more detail, RNAseq reads spanning the alternatively spliced exon junctions of *KRAS* were used to estimate the percentage of total *KRAS* transcripts made up by *KRAS4A* (see Methods). This analysis confirmed that *KRAS* mutant lung tumors on average have a significantly higher ratio of *KRAS4A* to *KRAS4B* than *KRAS* WT tumors. Similar results were recently reported by Stephens et al(*11*). To determine if splicing shifts towards higher *KRAS4A* expression in other cancer types, these analyses were extended to the TCGA pancreatic adenocarcinoma (PAAD) and colorectal adenocarcinoma (COADREAD) RNAseq datasets(*12*). Significant increases of *KRAS4A* and *KRAS4B* expression in *KRAS* mutant tumors were observed, with greater fold increase in *KRAS4A* (Fig. S5F). The percentage of total *KRAS* transcripts made up by *KRAS4A* showed similar increases in *KRAS* mutant PAAD and COADREAD, although significance was not reached, possibly due to the limited number of samples in these cohorts with known *KRAS* mutational status. Altogether, these data argue that the increase of *KRAS* expression observed in *KRAS* mutant cancers is driven not only by an overall increase in *KRAS* expression, but by altered splicing favoring an increase in the ratio of *KRAS4A* to *KRAS4B* transcripts.

### KRAS4A is enriched in cells with cancer stem cell properties

Tumors are highly heterogeneous, both at the cellular and genetic level, and sub-populations of tumor-initiating cells (also called cancer stem cells, or CSCs) exist that have metabolic requirements distinct from those of the bulk population of cancer cells(*13, 14*). The Kras4A isoform has been implicated in stem cell commitment during mouse development(*8*), raising the possibility that any differences between *KRAS4A* and *KRAS4B* knockout cells may reflect their expression in distinct subpopulations of cancer stem-or progenitor cells. We used a well characterized assay for “side population” cells(*15*) that have been shown to be enriched in stem cell properties, to investigate the expression of *KRAS4A* and *KRAS4B*. Expression of a marker of side population cells, (*ABCG2*)(*16*) was significantly elevated in side population cells from both A549 and SUIT2 cells, as well as from an additional pancreatic *KRAS* mutant cell line AsPC1 (Fig. 3A). Importantly, *KRAS4A* expression was also enriched in the same stem cell populations, as shown by specific TaqMan analysis of both *KRAS* splice isoforms (Fig. 3B). Homozygous knockout cells that no longer expressed KRAS4A had a reduced proportion of side population cells, which could be rescued by re-introduction of a functional *KRAS4A* expression construct (Figure 3C). The activity of the side population stem cell marker aldehyde dehydrogenase (ALDH)(*17*), was also reduced in the *KRAS4A* knockout cells, and restored by the re-introduced *KRAS4A* isoform(Fig.3C). In contrast to these observations regarding *KRAS4A*, there was no significant difference in *ALDH* expression in cells with reduced levels of KRAS4B in the heterozygous knockout lines (Fig. 3D) indicating that expression of *KRAS4A*, but not *KRAS4B* is linked to cells with stem cell properties.

**Figure 3.**
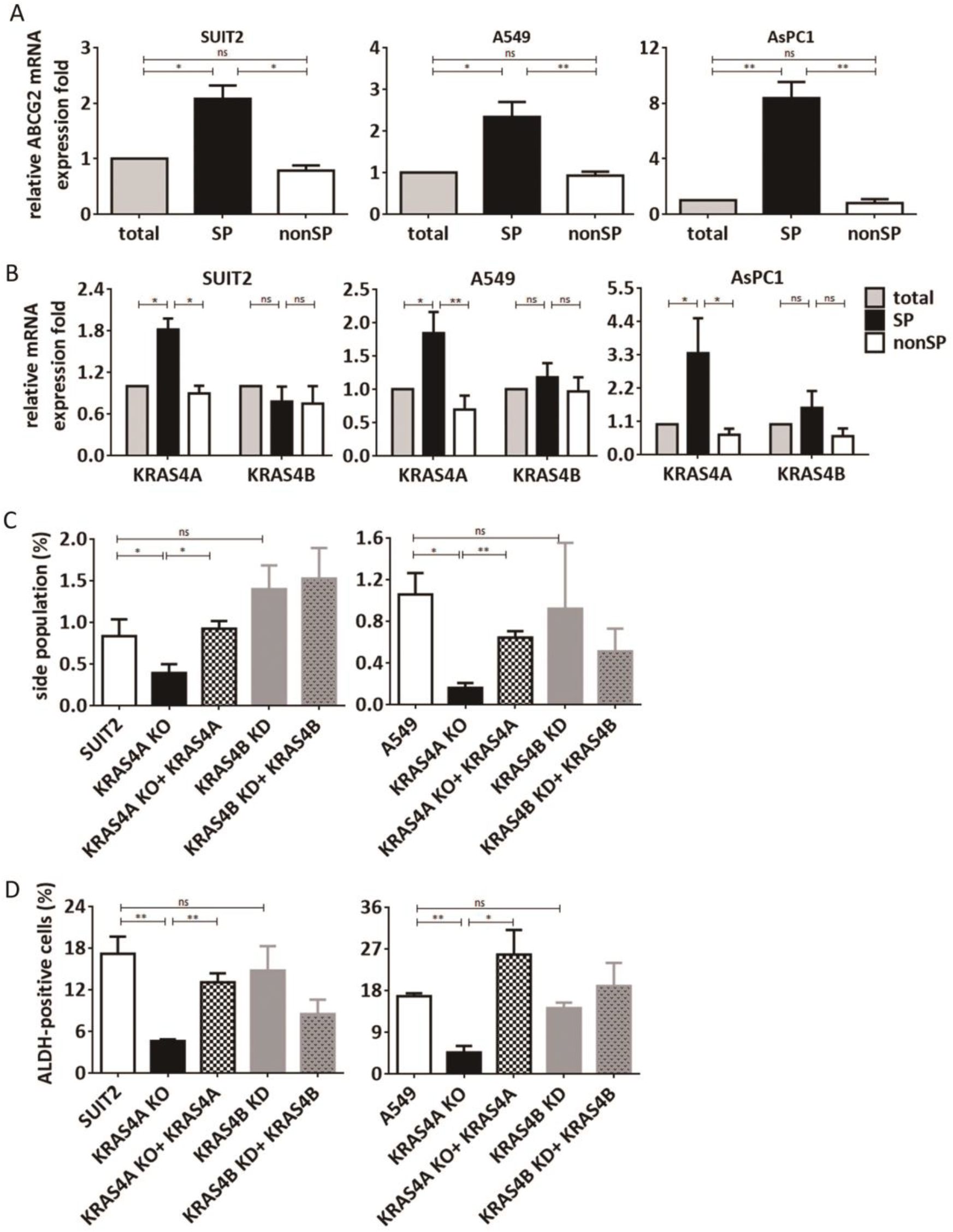
KRAS4A regulates stemness properties in human cancer cells. Stem cell side populations (SP) were assessed in A549, SUIT2 and AsPC1 cells by Hoechst 33342 staining. (A) *ABCG2* levels were significantly increased in side population cells from all 3 cell lines. (B) *KRAS4A*, but not *KRAS4B* was significantly increased in levels by TaqMan analysis of side population cells. Data are presented as mean ± s.e.m from three biological replicates. \**P* < 0.05; \*\**P* < 0.01 by one-way ANOVA with Tukey’s multiple comparison. (C-D) Loss of KRAS4A reduces the proportion of side population cells (C) as well as the expression of the ALDH side population marker (D). Both the proportion of side population cells and *ALDH* expression levels were restored by transfection of a *KRAS4A* expression construct using fluorescence-activated cell sorting analysis. Data are presented as mean ± s.e.m from three biological replicates. \**P* < 0.05; \*\**P* < 0.01 by Unpaired-t test.

### KRAS4A and 4B are induced by different types of cellular stress

Rapid tumor growth can lead to a hypoxic state which has been linked to stem cell activation and metabolic reprogramming(*18*). We investigated the possibility that hypoxia may specifically affect levels of *KRAS* splice variant expression in stem cells. Treatment of parental A549, SUIT2 and ASPC1 cells with CoCl_2_ induced an increase in expression of *HIF1A*, a known marker of hypoxia (Fig. 4A). *KRAS4A*, but not *KRAS4B* (Fig. 4B) showed a significant increase in expression as a consequence of this treatment, and the increase was predominantly in the side population cells rather than in the bulk cell population (Fig. 4C). In contrast, Endoplasmic Reticulum (ER) stress, which has been identified as a therapeutic target in *KRAS* mutant cancers(*19*) caused a significant increase in the ER marker *HSPA5* (also known as *GRP78*) and in *KRAS4B*, but not *KRAS4A*, in both A549 and SUIT2 cancer cell lines (Fig.4D, E). These data emphasize the functional differences between the two isoforms in the same human cancer cell lines.

**Figure 4:**
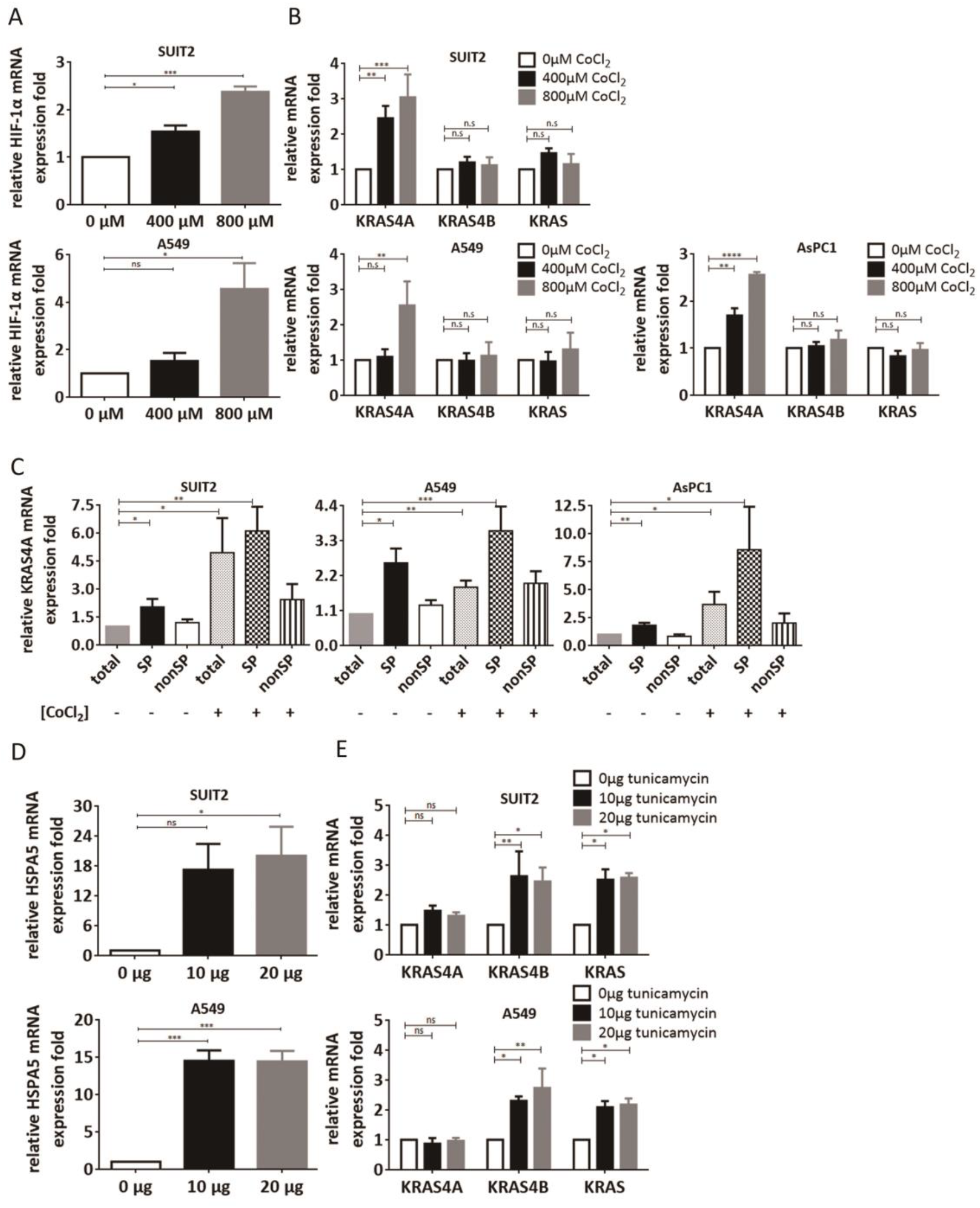
Different types of cellular stress affect expression of *KRAS* isoforms. (A-B) The *HIF-1α*, *KRAS4A*, *KRAS4B* and total *KRAS* mRNA levels were assessed by TaqMan analysis in A549, SUIT2 and AsPC1 cells after CoCl_2_ treatment for 48hr. The hypoxia marker *HIF-1α* was significantly increased by CoCl_2_ in all 3 cell lines. CoCl_2_ increased *KRAS4A*, but not *KRAS4B* levels in A549, SUIT2 and AsPC1 cells. Data are presented as mean ± s.e.m from three biological replicates. (C) The side populations were isolated with or without CoCl_2_ treatment and *KRAS4A* expression level was determined in sorted side population and non-side population cells by TaqMan analysis. *KRAS4A* was significantly increased by CoCl_2_ in side population cells. Data are presented as mean ± s.e.m from five biological replicates. (D-E) *HSPA5*, *KRAS4A*, *KRAS4B* and total *KRAS* mRNA level were assessed by TaqMan analysis in A549 and SUIT2 after tunicamycin treatment for 24hr. The ER stress marker *HSPA5* was significantly increased by tunicamycin. Tunicamycin increased *KRAS4B* and total *KRAS*, but not *KRAS4A* levels. Data are presented as mean ± s.e.m from three biological replicates. \**P* < 0.05; \*\**P* < 0.01; \*\*\**P* < 0.001 by one-way ANOVA with Tukey’s multiple comparison.

### KRAS4A and KRAS4B have different effects on tumor metabolism

Activation of *KRAS* has been linked to the increased glycolysis (Warburg Effect) seen in many cancer cell types(*20–22*). Although this was initially attributed to defective mitochondrial function, a more recent view is that mitochondrial oxidative phosphorylation is also important in tumors, and that coordination of these energy pathways is important for cancer progression(*23*). In fact, distinct metabolic pathways in CSC and non-CSC, have recently been demonstrated in several studies, suggesting models for coordinating energy requirements in stem and progenitor cells(*14, 24*). To investigate the effects of KRAS4A and KRAS4B on energy status, we first measured intracellular ATP levels after CRISPR-Cas9 knockout of *KRAS4A* and *KRAS4B* in SUIT2 and A549 cells. ATP content was significantly increased in the *KRAS4A* knockout cells, but was not changed by reduction of KRAS4B levels (Fig. 5A). The re-introduction of *KRAS4A* (but not a non-functional *KRAS4A* mutant (*7*) (see Methods) into *KRAS4A* knockout cells reduced ATP to levels similar to those in the parental cells (Fig. 5A). Increased lactate production was also seen in *KRAS4A* knockout cells, and this increase was rescued by expression of functional *KRAS4A* (Fig. 5B). In order to determine whether complete knockout of *Kras4A* or *Kras4B* in mice would have a similar effect, we analyzed levels of ATP and lactate in MEFs derived from *WT*, *Kras4A^−/−^* and *Kras4B^−/−^* mice. These showed similar changes to the human *KRAS4A* KO and *KRAS4B KD* cancer cells, respectively (Fig. 5A, B, right panels). In agreement with the observation of high ATP levels in human *KRAS4A^−/−^* cells, analysis of side population cells which are enriched in KRAS4A showed that these have reduced ATP levels (Fig. 5C)

**Figure. 5:**
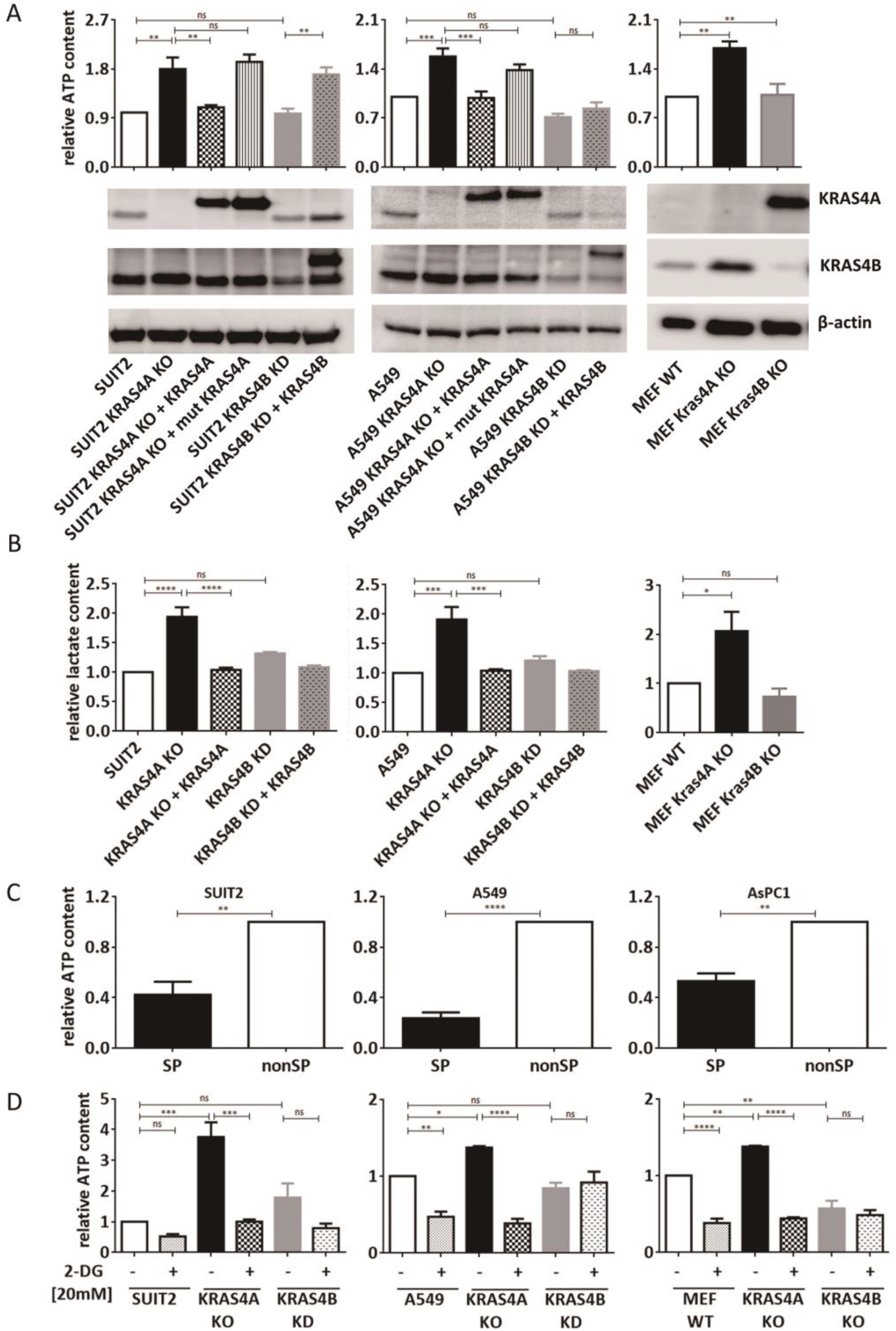
*KRAS* isoforms have different effects on energy status. (A) The ATP content of KRAS4A KO and KRAS4B KD cells (SUIT2, A549, and mouse MEFs) was assessed by bioluminescence assay. Compared to the corresponding parental cells (left and middle panels, Lanes 1), knockout of KRAS4A caused a significant increase in ATP levels (lanes 2) that was reduced by transfection of a functional *KRAS4A* construct (Lanes 3), but not by a *KRAS4A* construct mutated at the critical residue C180S(*7*) (lanes 4). Reduction of KRAS4B to levels that impacted tumor growth (Figure 3) did not cause an increase in ATP levels. Left panel: mouse MEFs homozygous for loss of *Kras4A* showed increased ATP levels but *Kras4B^−/−^* MEFs were similar to wild type cells. The expression of KRAS4A and KRAS4B protein was determined by Western blotting. (B) The intracellular lactate content in cell lysates was measured by lactate colorimetric assay kit, and showed a similar pattern to that seen for ATP, being higher in the KRAS4A knockout cells, both in human and mouse. Data are presented as mean ± s.e.m for four biological replicates. \*\**P* < 0.01; \*\*\**P* < 0.001 by one-way ANOVA with Tukey’s multiple comparison. (C) Side population cells were isolated by Hoechst 33342 staining and ATP content in sorted side population and non-side population cells was determined by bioluminescence assay. Side population cells have reduced levels of ATP. Data are presented as mean ± s.e.m for five biological replicates. \*\**P* < 0.01; \*\*\**P* < 0.001 by Unpaired-t test. (D) The glycolysis inhibitor, 2-DG, attenuated the increase of ATP content in KRAS4A stable knockout human cancer cells and mouse MEFs. Data are presented as mean ± s.e.m for three biological replicates. \**P* < 0.05; \*\**P* < 0.01; \*\*\**P* < 0.001; \*\*\*\**P* < 0.0001 by one-way ANOVA with Tukey’s multiple comparison.

To identify the mechanisms by which loss of KRAS4A leads to these metabolic changes, we treated *KRAS4A^−/−^* SUIT2, A549 and MEFs with the glycolysis inhibitor, 2-deoxyglucose (2-DG) (Fig. 5D). This treatment significantly decreased the intracellular ATP content for all cell types, and in particular reduced the high ATP levels in both human *KRAS4A^−/−^* cells and MEFs to parental control levels. In contrast, inhibitors of oxidative phosphorylation such as rotenone or oligomycin showed variable effects in different cell lines and no consistently significant reduction of ATP levels was seen in both *KRAS4A^−/−^* knockout cell lines (data not shown). Although the exact functions of KRAS4A and KRAS4B in tumor metabolism remain to be elucidated in detail, our data identify different consequences resulting from inactivation of each isoform, possibly reflecting the known heterogeneity in metabolic requirements in cancer stem and progenitor cells(*13, 14, 24, 25*). The relative levels of KRAS4A and KRAS4B in single cells may therefore impact metabolic changes required for the plasticity and interconversion capacity in stem and progenitor cell populations.

### Control of KRAS4A levels by the Rbm39/DCAF15 splice complex

Since *KRAS4A* mRNA levels were increased by hypoxia in the enriched stem cell population in 3 different *KRAS* mutant cell lines, but no effects were seen on *KRAS4B*, we reasoned that *KRAS* splicing may be differentially controlled in tumor stem and progenitor cells. Several small molecule inhibitors of different components of the splice site machinery have been identified, some of which can impact tumour cell growth(*26, 27*). We tested four of these inhibitors for effects on *KRAS4A/B* splicing: Pladienolide, which targets the splicing factor SF3B, disrupting the early stage of splice complex assembly(*28*); Isogenkgetin, a bioflavonoid and general splicing inhibitor(*29*); and two related sulfonamides Indisulam and Tasisulam, recently shown to inhibit the DCAF15/RBM39 splicing complex by inducing proteasomal degradation of the RBM39 RNA binding protein(*30, 31*). While Isogenkgetin had no obvious effect on levels of KRAS4A or KRAS4B mRNA, Pladienolide downregulated both KRAS4A and KRAS4B splice variants in A549 cells (Fig. 6A, B) in agreement with its general role in disruption of the splicing machinery. Treatment with Indisulam, or the related sulfonamide Tasisulam, specifically downregulated KRAS4A, but had no obvious effect on KRAS4B (Fig 6A, B). Similar results were obtained using two additional cell lines SUIT2 (Fig 6C, D), and AsPC1 (Fig.S6A, B), suggesting that the DCAF15/RBM39 complex is involved in control of KRAS4A levels. The downregulation of KRAS4A, but not KRAS4B, was also confirmed at the protein level in A549 and SUIT2 cells (Fig.6E, F), and in Aspc1 cells (Fig.S6C).

**Figure. 6:**
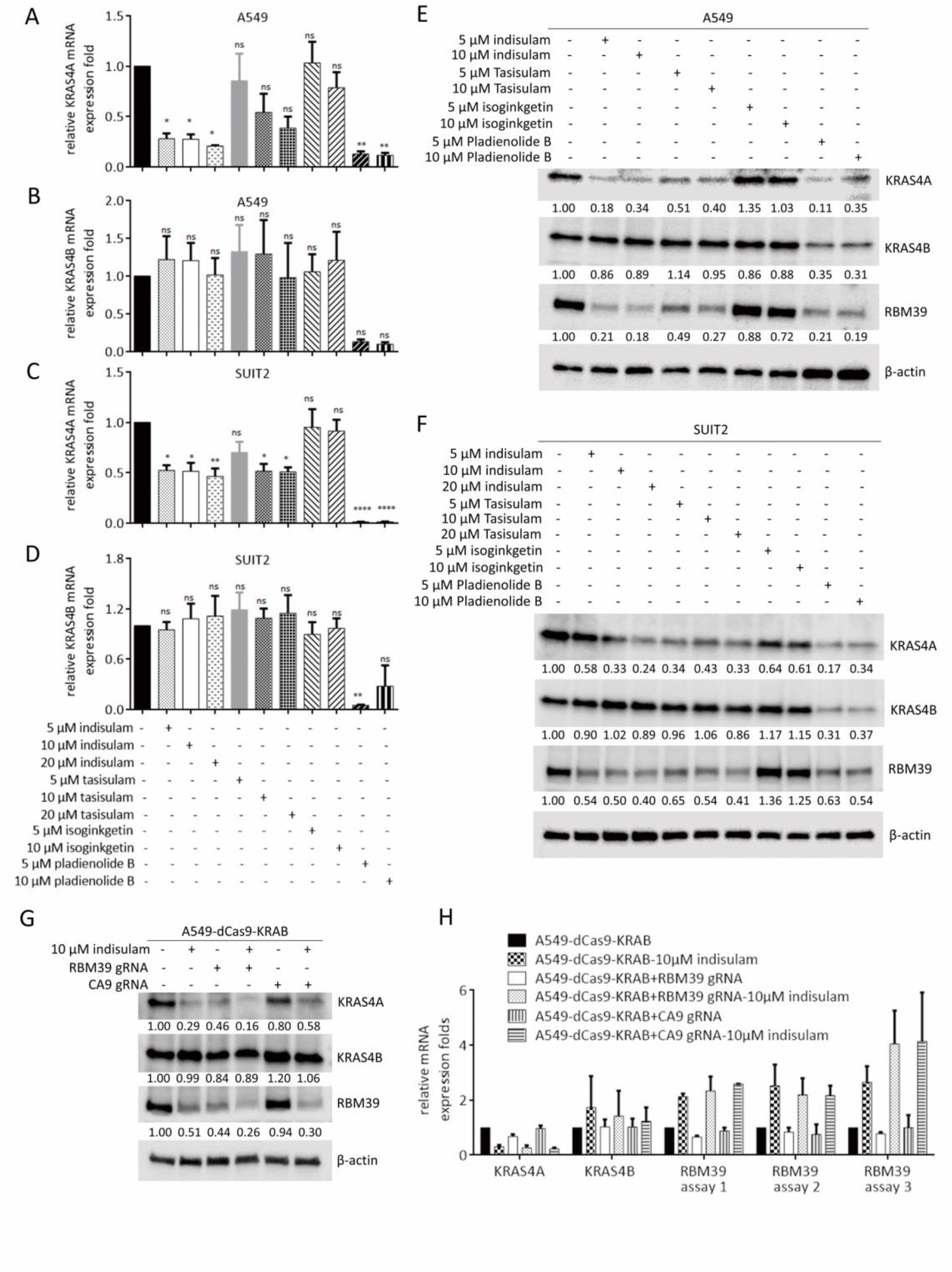
The RBM39 RNA-binding protein mediates *KRAS4A* splicing. (A-D) The *KRAS4A* and *KRAS4B* mRNA levels were assessed by TaqMan analysis in A549 (A and B) and SUIT2 (C and D) cells after small molecule inhibitor treatment for 48hr. The specific inhibitors used and their concentrations are shown below the plots. Data are presented as mean ± s.e.m for four biological replicates. \**P* < 0.05; \*\**P* < 0.01 by one-way ANOVA with Tukey’s multiple comparison. (E-F) The KRAS4A, KRAS4B and RBM39 protein levels were assessed in A549 (E) and SUIT2 (F) by Western blotting after small molecule inhibitor treatment for 48hr. Quantification of KRAS4A, KRAS4B and RBM39 levels was carried out using imageJ software. (G-H) Two sgRNAs targeting RBM39 or CA9 were transfected into BFP+A549 cells stably expressing dCas9-KRAB. Cells were incubated with small molecule inhibitors for 48hr and then analyzed by Western blotting (G) and Taqman analysis (H). The three Taqman assays (H) were carried out using probes for different *RBM39* exons.

To test the involvement of RBM39 in control of *KRAS* splicing, we first verified that Indisulam treatment indeed led to downregulation of RBM39 in A549 and SUIT2 cells (Fig 6E, F). In all cases where KRAS4A levels were reduced at RNA and protein levels, the RNA binding protein RBM39 was also reduced (Fig. 6E, F and Fig. S6C). This effect was seen in all three cell lines for both Indisulam and Tasisulam, which target RBM39 through its interaction with DCAF15(*30*) but also with the general splice inhibitor Pladienolide, although the molecular basis for this latter observation is unclear.

We then generated RBM39 knockdown cells using CRISPRi/dCas9-KRAB(*32*) using specific guide RNAs targeting the *RBM39* gene in A549 cells. Depletion of RBM39 caused a significant reduction in KRAS4A splice variant level but had no effect on KRAS4B (Fig. 6G), attesting to the specificity of RBM39 in control of *KRAS* pre-mRNA splicing. Although the RBM39 guide RNAs had a significant effect on RBM39 protein in A549 cells (Fig. 6G), the effect at the mRNA level was more modest (Fig. 6H). It is possible that this reflects a feedback activation of *RBM39* transcription when the protein is depleted, as Indisulam treatment surprisingly caused a significant upregulation in RBM39 mRNA while decreasing the protein levels (Fig. 6G, H). Further studies of this apparent feedback control of RBM39, which seems variable between cell lines (data not shown) may shed light on the exact mechanisms involved.

Indisulam was originally proposed to act as an inhibitor of CA9 (carbonic anhydrase 9) which is involved in HIF1A regulation and hypoxia(*33*). However CRISPRi/dCas9-KRAB downregulation of CA9 mRNA (Fig. S6D) had no effect on levels of KRAS4A, or on the impact of Indisulam on KRAS4A levels (Fig 6G). While RBM39 influences splicing of many pre-mRNAs genome-wide, these data support the proposal(*30*) that RBM39 has a more limited set of physiological targets than would be expected for a general inhibitor of the splice machinery, and identify *KRAS4A* as a specific RBM39 target that can be inhibited by the small molecule drug Indisulam.

## Discussion

### Essential functions for both KRAS4A and KRAS4B in cancer development

The existence of two distinct isoforms of *Kras* has been known for many years, but most research has focused on the *KRAS4B* isoform which is widely and abundantly expressed across a range of tissues. Both isoforms are required for initiation and/or progression of carcinogenesis in the lung, as tumors are only induced in animals that have at least one functional *Kras* allele capable of expressing both proteins. In spite of the fact that mutant forms of either Kras4A or Kras4B can transform cells in culture(*34, 35*) mutation of either endogenous isoform alone is insufficient to initiate carcinogenesis. This resistance is not due to loss of accessibility to the initiating mutagen, as *Hras*, when inserted into the same locus under the control of the Kras regulatory elements in exactly the same way, can be mutated by urethane, giving rise to highly aggressive lung cancers(*5*). Double heterozygous *Kras4A^+/−^/Kras4B^+/−^* mice were also extremely resistant, even under conditions where the carcinogen dose was significantly increased, suggesting that coordinated splicing to generate mutant versions of both isoforms from the same allele is required for tumorigenesis.

*Kras4A* is expressed during differentiation of pluripotent embryonic stem cells and in a subset of cells in adult tissues(*8*), raising the possibility that Kras4A has specific functions in a small population of cells with stem cell properties(*5*), whereas Kras4B protein is more widely expressed in both stem and progenitor cells. Here we have shown that expression of *KRAS4A*, but not *KRAS4B*, is enriched in stem cell-like side population cells from human *KRAS* mutant tumours. A number of complementary experiments support the conclusion that KRAS4A plays a specific role in this subset of enriched stem cells: 1) Loss of KRAS4A, but not KRAS4B, causes a decrease in the proportion of cells with side population characteristics, as well as decreased activity of ALDH, a known marker of these cells; 2) hypoxic conditions which are known to lead to reactivation of stem cells, cause an upregulation of *KRAS4A*, but not *KRAS4B*; 3) Loss of *KRAS4A* but not *KRAS4B* induced a significant increase in ATP and lactate levels, particularly in the side population stem-like cells. In contrast, *KRAS4B* was equally expressed in both side population and non-side population cells, and showed specific induction by exposure to ER stress. We propose that the splicing of *KRAS* to generate the 4A and 4B isoforms may be a critical event in controlling the metabolic requirements in stem and progenitor cells, as well as in orchestrating the responses of these different cells to hypoxia or ER stress (Fig.7). The transition from cancer stem to progenitor cells is reversible depending on the tumor microenvironment, and can be influenced by inflammation or stromal components within tumors(*36*). This plasticity may be facilitated by splicing of *KRAS* to generate the 4A and 4B isoforms, with their complementary functions in coordinating the stresses associated with rapid tumor growth. Alternative splicing has also evolved as a mediator of the IRE1 arm of the Unfolded Protein Response (UPR) by activating splicing of the short form of XBP1 to modulate downstream protein synthesis and accumulation in the ER(*37*). The balance between KRAS4A and 4B splice variants may also play a more general role in responding to the stresses of tissue regeneration and tumor growth.

**Figure 7.**
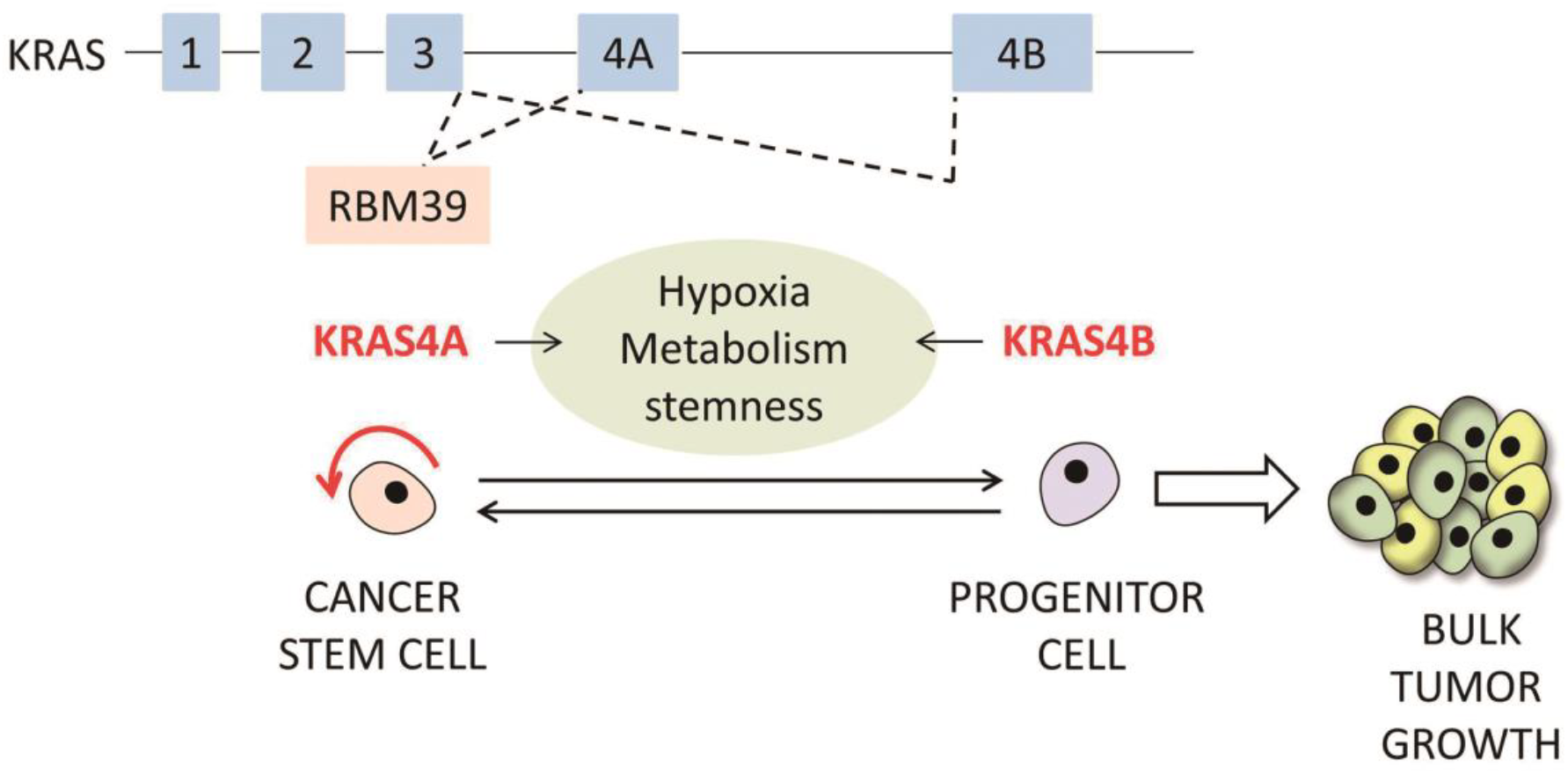
Model for control of stem-progenitor cell transition by *KRAS4A/4B* splicing. KRAS4A is enriched in the cancer stem cell population, whereas KRAS4B is more ubiquitously expressed in stem and progenitor cells. Regulation of the expression of *KRAS4A*, in part through the DCAF15/RBM39 splicing complex, helps to modulate the metabolic requirements and stress responses associated with the cancer stem-progenitor cell transition.

Our identification of Indisulam as an inhibitor of *KRAS4A* splicing, which has no obvious effect on *KRAS4B*, provides a novel route to direct targeting of one of the mutant *KRAS* splice isoforms. Further studies may reveal vulnerabilities in cells in which KRAS4A has been depleted or pharmacologically downregulated using Indisulam or other small molecule drugs that impact this pathway. Furthermore, mutations in *KRAS* and/or levels of mutant KRAS4A may provide biomarkers for sensitivity to the effects of Indisulam or other splice regulators. The availability of mouse models and human cells that exclusively or preferentially express one splice variant will provide us with new opportunities to identify isoform-specific vulnerabilities to treatment with inhibitors of the RAS or other signaling pathways, leading to effective combinatorial treatments for *KRAS* mutant tumors.

## Methods

### Mice

The Kras4B KO mice were generated as previously described(*4*) and chemical carcinogenesis of lung was performed by intraperitoneal injection of urethane as previously described(*5*). Mice were injected at approximately 8-10 weeks of age, sacrificed 20 weeks later, and tumor size and number assessed under a dissecting microscope with the aid of a ruler and reference images of a range of circles with different diameters. All tumor scoring was performed blind to mouse genotype.

### Cell culture

SUIT2, A549 and mouse embryonic fibroblasts (MEFs) isolated from E13.5 embryos were cultured in high glucose DME media (Gibco) containing 10% fetal bovine serum (Gibco) and 1% penicilin-streptamicine (Gibco).

### Generation of KRAS4A, KRAS4B and RBM39 knockout or knockdown cells

Human KRAS4A knockout and KRAS4B knockdown cells were generated by CRISPR-Cas9-mediated genome engineering as previously described(*38*). sgRNA targets were GGAGGATGCTTTTTATACAT for KRAS4A and TTCTCGAACTAATGTATAGA for KRAS4B. To confirm the insertions and deletions of each allele, the PCR products around the CRISPR-cas9 targeted sites were amplified from gDNA of established stable clones and cloned into pMiniT 2.0 vector (NEB), followed by plasmid DNA isolation and sanger sequencing. The primers used were caaaccaggattctagcccata and gtggttgccaccttgttacc for KRAS4A and ttcagttgcctgaagagaaaca and agtctgcatggagcaggaaa for KRAS4B. Human RBM39 and CA9 knockdown cells were generated by CRISPRi/dCas9-KRAB-mediated genome engineering as previously described(*32*). A549 cells stably expressing dCas9-KRAB were enriched by flow cytometry for BFP expression and two sgRNA targeting RBM39 or CA9 were transfected into A549-dCas9-KRAB cell for puromycin selection. sgRNA targets were GAGCAGCGGCCGCCATTTCA and GGAGAGCAGGACGGCGGCTT for RBM39 and GGGATCAACAGAGGGAGCCA and GCAGGGGCCGGGATCAACAG for CA9. The shRNA-mediated knockdown cells were generated by antibiotic selection after infection with retroviral particles collected in supernatants of Phoenix cells transfected with pSuper.retro plasmid carrying the shRNA sequences. shRNA targets GGTGAGGGAGATCCGACAATA for KRAS4A and GACAGGGTGTTGATGATGCCT for KRAS4B.

### Western Blotting analysis

Cells were lysed with RIPA buffer (Thermo Scientific) and lysates concentrations were determined by BCA protein assay (Thermo Scientific). 80ug lysates were subjected to 4-12% SDS-PAGE (Bio-Rad) and then transferred to PVDF membrane. PVDF membranes were blocked in Tris-buffered saline 0.1% Tween-20 (TBST) containing 5% non-fat milk for 1h at room temperature and incubated overnight with primary antibody diluted in TBST containing 3% non-fat milk at 4°C. Membranes were washed with TBST and incubated with horseradish peroxidase (HRP)-conjugated secondary antibodies diluted in TBST containing 3% non-fat milk at 4°C for 2h. The primary antibodies used were Hras (sc520; Santa Cruz), Nras (sc519; Santa Cruz), phospho-p44/p42 Map Kinase, phospho-Akt, phospho-Erk, β-actin (sc47778; Santa Cruz); KRAS4B (WH0003845M1; sigma); rat anti-KRAS4A (10C11E4, custom antibody) and RBM39 (WH0009584M1; sigma). Proteins were visualized using the enhanced chemiluminescence (ECL) system (ECL™ Prime Western Blotting System, GE Healthcare Bioscience).

### Antibody generation

Custom rat anti-KRAS4A antibody was developed by Genscript using the peptide sequence CEIRQYRLKKISKEEK as antigen for immunization..

### Cell proliferation assay

1,000 cells were plated in 96-well culture plates and cell proliferation was determined by CyQUANT cell proliferation assay kit (C35011, Invitrogen) as described by the manufacturer.

### Colony formation

Cells were plated at density of 500 cells per well in six-well plate and incubated for 10 days at 37°C in DMEM medium containing 10% fetal bovine serum and 1% penicilin-streptamycin. Colonies were counted after methanol fixation and crystal violet staining.

### Soft agar growth assays

5,000 cells were mixed with 1 ml of 0.3% agar in DMEM supplemented with 10% FBS and layered onto 1.5 ml 0.6% agar in DMEM supplemented with 10% FBS. The colonies were incubated for 2 weeks in SUIT2 cells and for 3 weeks in A549 cells. At the end of incubation, the colonies were stained with crystal violet and the images were captured using microscope followed by quantification of colony area using Image-Pro Plus software.

### Xenograft tumor model

SUIT2 cells (4 × 10^6^) and A549 cells (3 × 10^6^) were dispersed in 75 μL DMEM and 75 μL Matrigel (356230, BD Biosciences) and injected subcutaneously into nude mice. Tumor volume was calculated as follows: V = L × W^2^ × 0.52, where L and W represent length and width respectively.

### Site-directed mutagenesis

Site-directed mutagenesis to generate the mutant RAS clones was performed with a QuikChange mutagenesis kit (Stratagene) and the primers for amino acid 180 site mutation of KRAS4A are CAGCAAAGAAGAAAAGACTCCTGGCAGTGTGAAAATT and AATTTTCACACTGCCAGGAGTCTTTTCTTCTTTGCTG.

### Measurement of intracellular ATP

5 × 10^5^ cells were collected 48 hours after cell seeding and were lysed with 0.1ml reaction assay buffer supplied in the ATP detection kit (A22066, Thermo Fischer Scientific) followed by centrifugation at 16,000 x g for 5 min. The cellular ATP levels in the supernatants were determined according to the instructions of the manufacturer. For the ATP depletion analysis, cells were incubated with 20mM 2DG (D6134, sigma), 1μM oligomycin (495455, Calbiochem) and 1μM rotenone for (R8875, sigma) 24 h followed by ATP level determination.

### Measurement of intracellular lactate

3 × 10^6^ cells were collected 48 hours after cell seeding and were lysed with 0.1 ml lactate assay buffer supplied in the L-Lactate assay kit (ab65330, Abcam) followed by centrifugation at 16,000 x g for 10 min. The supernatants were processed to remove the endogenous LDH by deproteinizing sample preparation Kit (ab204708, Abcam). The intracellular lactate concentration was determined on deproteinized supernatant samples by L-Lactate assay kit according to the manufacturer’s instructions

### Flow cytometry

For side population identification and isolation analysis, cells were suspended at 1×10^6^ cells per ml in DMEM containing 2% FBS, 10mM HEPES and 5μg/ml Hoechst33342 dye (B2261, Sigma Aldrich), either alone or combination with 50μM verapamil (ALX-550-306-G001, Enzo Life Sciences) and incubated in water bath at 37 °C for 2h. After incubation, cells were suspended in PBS containing 2% FBS and 10mM HEPES on ice and stained with 1μg/ml propidium iodide followed by sorting or analysis using AriaII (BD) fluorescence activated cell sorting system (FACS). For the ALDH activity analysis, cells were suspended at 1×10^6^ cells per ml in ALDEFLUOR assay buffer containing activated ALDEFLUOR™ Reagent, either alone or combination with diethyl aminobenzaldehyde (DEAB), and incubated 30 min at 37 °C according to the manufacturer’s instruction in ALDEFLUOR kit (01700, StemCell Technologies). For the glucose uptake analysis, cells were suspended at 1.5×10^6^ cells per ml in FBS containing 1% FBS, 5mM HEPES and 2‐ ‐(7‐nitrobenz‐2‐oxa‐1,3‐diazol‐4‐yl)amino]‐2‐ deoxyglucose (2-NBDG, N13195, invitrogen) 4μM for SUIT2 and 8μM for A549 and incubated in water bath at 37 °C for 15 min, followed by FACS analysis.

### RNA extraction and quantitative polymerase chain reaction (qPCR)

RNA was extracted using TRIzol Reagent (15596026, Invitrogen), and cDNA was synthesized using iScript Synthesis kit (1708840, Bio-Rad) according to the manufacturer’s instructions. Quantitative real-time RT-PCR was carried out using Taqman Mix in an ABI Prism7900HT Sequence Detection System (Applied Biosystems, Foster City, CA). TaqMan Gene expression assays used were as follows: KRAS (Hs00364282_m1), KRAS4B (Hs00270666_m1 KRAS), HPRT1 (Hs02800695_m1) and ABCG2 (Hs01053790_m1). The primers and probe used for amplification of KRAS4A were as follows: TGTGATTTGCCTTCTAGAACAGTAGAC, CTCACCAATGTATAAAAAGCATCCTC, and 5’-FAM-CGAAACAGGCTCAGGAG-MGB-3’. Relative mRNA expression levels were normalized to HPRT1.

### TCGA analysis of *KRAS4A* and *KRAS4B* isoform-specific expression

TCGA lung adenocarcinoma (LUAD), pancreatic adenocarcinoma (PAAD), and colorectal adenocarcinoma (COADREAD) RNA sequencing and clinical annotation data were downloaded from UCSC Cancer Genome Browser (now UCSC Xena). Specifically, level 3 normalized RSEM values for reads spanning splice junctions was downloaded, and used to calculate frequencies of reads spanning junctions of exons 4A and 4B of *KRAS*. Given that splicing to yield *KRAS4B* results in exclusion of exon 4A, while *KRAS4A* results in inclusion of both exons 4A and 4B, an exon 4A inclusion score was calculated to determine the fraction of *KRAS* transcripts made up by *KRAS4A*. This score is calculated as ((inc1 + inc2)/2) / (((inc1 + inc2)/2) + exc), where inc1 and inc2 are reads spanning the junction of exons 3 and 4A, and reads spanning the junction of exons 4A and 4B, respectively, and exc are reads spanning the junction of exons 3 and 4B. To prevent artificially low scores in samples with lower overall *KRAS* expression or read coverage, particularly those that are *KRAS* WT, samples with less than 20 exc reads were not included in the analysis. No limit was set on inc1 and inc2 reads, as it is biologically plausible that some samples have minimal *KRAS4A* expression.

### Data analysis and generation of plots

Data were analyzed and non-parametric statistical tests performed in R. Plots were generated using the R package ggplot2 (H. Wickham. ggplot2: Elegant Graphics for Data Analysis. Springer-Verlag New York, 2009.)

## Acknowledgements

We are greatly appreciative of help and comments from our colleagues in refining this study and manuscript.

## Funding

This work was supported by US National Cancer Institute (NCI) grants RO1CA184510, UO1 CA176287, and R35CA210018 and the Barbara Bass Bakar Professorship of Cancer Genetics. Wei-Ching Chen was supported by a Fellowship from Taiwan Ministry of Science and Technology and by the UCSF Pancreas Center and the Schwartz Family Foundation. P.M.K.W was supported by NIH training grant T32 GM007175 and an NCI F31 NRSA award. MDT was supported by a UCSF Senate Research grant.

## Author contributions

W-C.C designed and carried out most of the biochemical experiments shown in Figs 2-6 for characterization of Kras4A and Kras4B cells, and generated Crispri/dCas9-KRAB knockdown cells, MD To helped with the overall study design, generated the Kras4BKO mouse, carried out shRNA studies and mouse carcinogenesis experiments, PW generated and characterized Crispr/Cas9 knockout cells and carried out bioinformatics analysis, animal carcinogenesis studies and tumor analysis, MP provided guidance and protocols for biochemical and metabolic assays and helped with data analysis and interpretation, RD performed mouse carcinogenesis studies, IJ K carried out TaqMan analysis of *KRAS4A/B* expression in human tumours, QT performed mouse tumour mutation analysis and helped with cell culture experiments, NB characterized Crispr/Cas9 knockout cells, AB designed and supervised the overall study. The paper was written by AB, MDT, WCC, and PW, with contributions from the other authors.

## Competing Interests

Authors declare no competing interests.

## Data and Materials availability

All data is available in the main text or the supplementary materials. All knockout mouse strains and cell lines are available on request upon signing the appropriate UCSF Materials Transfer Agreement.

## References

1. J. F. Hancock, Ras proteins: different signals from different locations. Nature reviews. Molecular cell biology 4, 373–384 (2003); published online EpubMay (10.1038/nrm1105).

2. L. Johnson, D. Greenbaum, K. Cichowski, K. Mercer, E. Murphy, E. Schmitt, R. T. Bronson, H. Umanoff, W. Edelmann, R. Kucherlapati, T. Jacks, K-ras is an essential gene in the mouse with partial functional overlap with N-ras. Genes Dev 11, 2468–2481 (1997); published online EpubOct 1 (

3. S. J. Plowman, D. J. Williamson, M. J. O’Sullivan, J. Doig, A. M. Ritchie, D. J. Harrison, D. W. Melton, M. J. Arends, M. L. Hooper, C. E. Patek, While K-ras is essential for mouse development, expression of the K-ras 4A splice variant is dispensable. Mol Cell Biol 23, 9245–9250 (2003); published online EpubDec (

4. N. Potenza, C. Vecchione, A. Notte, A. De Rienzo, A. Rosica, L. Bauer, A. Affuso, M. De Felice, T. Russo, R. Poulet, G. Cifelli, G. De Vita, G. Lembo, R. Di Lauro, Replacement of K-Ras with H-Ras supports normal embryonic development despite inducing cardiovascular pathology in adult mice. EMBO reports 6, 432–437 (2005); published online EpubMay (10.1038/sj.embor.7400397).

5. M. D. To, C. E. Wong, A. N. Karnezis, R. Del Rosario, R. Di Lauro, A. Balmain, Kras regulatory elements and exon 4A determine mutation specificity in lung cancer. Nature genetics 40, 1240–1244 (2008); published online EpubOct (10.1038/ng.211).

6. C. E. Patek, M. J. Arends, W. A. Wallace, F. Luo, S. Hagan, D. G. Brownstein, L. Rose, P. S. Devenney, M. Walker, S. J. Plowman, R. L. Berry, W. Kolch, O. J. Sansom, D. J. Harrison, M. L. Hooper, Mutationally activated K-ras 4A and 4B both mediate lung carcinogenesis. Experimental cell research 314, 1105–1114 (2008); published online EpubMar 10 (10.1016/j.yexcr.2007.11.004).

7. F. D. Tsai, M. S. Lopes, M. Zhou, H. Court, O. Ponce, J. J. Fiordalisi, J. J. Gierut, A. D. Cox, K. M. Haigis, M. R. Philips, K-Ras4A splice variant is widely expressed in cancer and uses a hybrid membrane-targeting motif. Proceedings of the National Academy of Sciences of the United States of America 112, 779–784 (2015); published online EpubJan 20 (10.1073/pnas.1412811112).

8. S. Pells, M. Divjak, P. Romanowski, H. Impey, N. J. Hawkins, A. R. Clarke, M. L. Hooper, D. J. Williamson, Developmentally-regulated expression of murine K-ras isoforms. Oncogene 15, 1781–1786 (1997); published online EpubOct 9 (10.1038/sj.onc.1201354).

9. Comprehensive molecular profiling of lung adenocarcinoma. Nature 511, 543–550 (2014); published online EpubJul 31 (10.1038/nature13385).

10. I. J. Kim, D. Quigley, M. D. To, P. Pham, K. Lin, B. Jo, K. Y. Jen, D. Raz, J. Kim, J. H. Mao, D. Jablons, A. Balmain, Rewiring of human lung cell lineage and mitotic networks in lung adenocarcinomas. Nature communications 4, 1701 (2013)10.1038/ncomms2660).

11. R. M. Stephens, M. Yi, B. Kessing, D. V. Nissley, F. McCormick, Tumor RAS Gene Expression Levels Are Influenced by the Mutational Status of RAS Genes and Both Upstream and Downstream RAS Pathway Genes. Cancer informatics 16, 1176935117711944 (2017)10.1177/1176935117711944).

12. Comprehensive molecular characterization of human colon and rectal cancer. Nature 487, 330–337 (2012); published online EpubJul 18 (10.1038/nature11252).

13. A. Viale, P. Pettazzoni, C. A. Lyssiotis, H. Ying, N. Sanchez, M. Marchesini, A. Carugo, T. Green, S. Seth, V. Giuliani, M. Kost-Alimova, F. Muller, S. Colla, L. Nezi, G. Genovese, A. K. Deem, A. Kapoor, W. Yao, E. Brunetto, Y. Kang, M. Yuan, J. M. Asara, Y. A. Wang, T. P. Heffernan, A. C. Kimmelman, H. Wang, J. B. Fleming, L. C. Cantley, R. A. DePinho, G. F. Draetta, Oncogene ablation-resistant pancreatic cancer cells depend on mitochondrial function. Nature 514, 628–632 (2014); published online EpubOct 30 (10.1038/nature13611).

14. A. Viale, G. F. Draetta, Metabolic Features of Cancer Treatment Resistance. Recent Results Cancer Res. 207, 135–156 (2016)10.1007/978-3-319-42118-6_6).

15. S. Orecchioni, F. Bertolini, Characterization of Cancer Stem Cells. Methods Mol Biol 1464, 49–62 (2016)10.1007/978-1-4939-3999-2_5).

16. M. M. Ho, A. V. Ng, S. Lam, J. Y. Hung, Side population in human lung cancer cell lines and tumors is enriched with stem-like cancer cells. Cancer research 67, 4827–4833 (2007); published online EpubMay 15 (10.1158/0008-5472.CAN-06-3557).

17. F. Jiang, Q. Qiu, A. Khanna, N. W. Todd, J. Deepak, L. Xing, H. Wang, Z. Liu, Y. Su, S. A. Stass, R. L. Katz, Aldehyde dehydrogenase 1 is a tumor stem cell-associated marker in lung cancer. Molecular cancer research: MCR 7, 330–338 (2009); published online EpubMar (10.1158/1541-7786.MCR-08-0393).

18. E. B. Rankin, J. M. Nam, A. J. Giaccia, Hypoxia: Signaling the Metastatic Cascade. Trends in cancer 2, 295–304 (2016); published online EpubJun (10.1016/j.trecan.2016.05.006).

19. T. De Raedt, Z. Walton, J. L. Yecies, D. Li, Y. Chen, C. F. Malone, O. Maertens, S. M. Jeong, R. T. Bronson, V. Lebleu, R. Kalluri, E. Normant, M. C. Haigis, B. D. Manning, K. K. Wong, K. F. Macleod, K. Cichowski, Exploiting cancer cell vulnerabilities to develop a combination therapy for ras-driven tumors. Cancer cell 20, 400–413 (2011); published online EpubSep 13 (10.1016/j.ccr.2011.08.014).

20. R. Serna-Blasco, M. Sanz-Alvarez, O. Aguilera, J. Garcia-Foncillas, Targeting the RAS-dependent chemoresistance: The Warburg connection. Semin. Cancer Biol., (2018); published online EpubFeb 9 (10.1016/j.semcancer.2018.01.016).

21. E. M. Kerr, E. Gaude, F. K. Turrell, C. Frezza, C. P. Martins, Mutant Kras copy number defines metabolic reprogramming and therapeutic susceptibilities. Nature 531, 110–113 (2016); published online EpubMar 3 (10.1038/nature16967).

22. H. Ying, A. C. Kimmelman, C. A. Lyssiotis, S. Hua, G. C. Chu, E. Fletcher-Sananikone, J. W. Locasale, J. Son, H. Zhang, J. L. Coloff, H. Yan, W. Wang, S. Chen, A. Viale, H. Zheng, J. H. Paik, C. Lim, A. R. Guimaraes, E. S. Martin, J. Chang, A. F. Hezel, S. R. Perry, J. Hu, B. Gan, Y. Xiao, J. M. Asara, R. Weissleder, Y. A. Wang, L. Chin, L. C. Cantley, R. A. DePinho, Oncogenic Kras maintains pancreatic tumors through regulation of anabolic glucose metabolism. Cell 149, 656–670 (2012); published online EpubApr 27 (10.1016/j.cell.2012.01.058).

23. N. Hay, Reprogramming glucose metabolism in cancer: can it be exploited for cancer therapy? Nature reviews. Cancer 16, 635–649 (2016); published online EpubOct (10.1038/nrc.2016.77).

24. P. Sancho, E. Burgos-Ramos, A. Tavera, T. Bou Kheir, P. Jagust, M. Schoenhals, D. Barneda, K. Sellers, R. Campos-Olivas, O. Grana, C. R. Viera, M. Yuneva, B. Sainz, Jr., C. Heeschen, MYC/PGC-1alpha Balance Determines the Metabolic Phenotype and Plasticity of Pancreatic Cancer Stem Cells. Cell metabolism 22, 590–605 (2015); published online EpubOct 6 (10.1016/j.cmet.2015.08.015).

25. S. Shibao, N. Minami, N. Koike, N. Fukui, K. Yoshida, H. Saya, O. Sampetrean, Metabolic heterogeneity and plasticity of glioma stem cells in a mouse glioblastoma model. Neuro-oncology 20, 343–354 (2018); published online EpubFeb 19 (10.1093/neuonc/nox170).

26. S. Bonnal, L. Vigevani, J. Valcarcel, The spliceosome as a target of novel antitumour drugs. Nature reviews. Drug discovery 11, 847–859 (2012); published online EpubNov (10.1038/nrd3823).

27. R. Assi, H. M. Kantarjian, T. M. Kadia, N. Pemmaraju, E. Jabbour, N. Jain, N. Daver, Z. Estrov, T. Uehara, T. Owa, J. E. Cortes, G. Borthakur, Final results of a phase 2, open-label study of indisulam, idarubicin, and cytarabine in patients with relapsed or refractory acute myeloid leukemia and high-risk myelodysplastic syndrome. Cancer 124, 2758–2765 (2018); published online EpubJul 1 (10.1002/cncr.31398).

28. Y. Kotake, K. Sagane, T. Owa, Y. Mimori-Kiyosue, H. Shimizu, M. Uesugi, Y. Ishihama, M. Iwata, Y. Mizui, Splicing factor SF3b as a target of the antitumor natural product pladienolide. Nature chemical biology 3, 570–575 (2007); published online EpubSep (10.1038/nchembio.2007.16).

29. K. O’Brien, A. J. Matlin, A. M. Lowell, M. J. Moore, The biflavonoid isoginkgetin is a general inhibitor of Pre-mRNA splicing. The Journal of biological chemistry 283, 33147–33154 (2008); published online EpubNov 28 (10.1074/jbc.M805556200).

30. T. Han, M. Goralski, N. Gaskill, E. Capota, J. Kim, T. C. Ting, Y. Xie, N. S. Williams, D. Nijhawan, Anticancer sulfonamides target splicing by inducing RBM39 degradation via recruitment to DCAF15. Science 356, (2017); published online EpubApr 28 (10.1126/science.aal3755).

31. T. Uehara, Y. Minoshima, K. Sagane, N. H. Sugi, K. O. Mitsuhashi, N. Yamamoto, H. Kamiyama, K. Takahashi, Y. Kotake, M. Uesugi, A. Yokoi, A. Inoue, T. Yoshida, M. Mabuchi, A. Tanaka, T. Owa, Selective degradation of splicing factor CAPERalpha by anticancer sulfonamides. Nature chemical biology 13, 675–680 (2017); published online EpubJun (10.1038/nchembio.2363).

32. L. A. Gilbert, M. H. Larson, L. Morsut, Z. Liu, G. A. Brar, S. E. Torres, N. Stern-Ginossar, O. Brandman, E. H. Whitehead, J. A. Doudna, W. A. Lim, J. S. Weissman, L. S. Qi, CRISPR-mediated modular RNA-guided regulation of transcription in eukaryotes. Cell 154, 442–451 (2013); published online EpubJul 18 (10.1016/j.cell.2013.06.044).

33. F. Abbate, A. Casini, T. Owa, A. Scozzafava, C. T. Supuran, Carbonic anhydrase inhibitors: E7070, a sulfonamide anticancer agent, potently inhibits cytosolic isozymes I and II, and transmembrane, tumor-associated isozyme IX. Bioorganic & medicinal chemistry letters 14, 217–223 (2004); published online EpubJan 5 (

34. K. Shimizu, D. Birnbaum, M. A. Ruley, O. Fasano, Y. Suard, L. Edlund, E. Taparowsky, M. Goldfarb, M. Wigler, Structure of the Ki-ras gene of the human lung carcinoma cell line Calu-1. Nature 304, 497–500 (1983); published online EpubAug 11–17 (

35. J. K. Voice, R. L. Klemke, A. Le, J. H. Jackson, Four human ras homologs differ in their abilities to activate Raf-1, induce transformation, and stimulate cell motility. The Journal of biological chemistry 274, 17164–17170 (1999); published online EpubJun 11 (

36. M. Holzel, A. Bovier, T. Tuting, Plasticity of tumour and immune cells: a source of heterogeneity and a cause for therapy resistance? Nature reviews. Cancer 13, 365–376 (2013); published online EpubMay (10.1038/nrc3498).

37. P. Walter, D. Ron, The unfolded protein response: from stress pathway to homeostatic regulation. Science 334, 1081–1086 (2011); published online EpubNov 25 (10.1126/science.1209038).

38. F. A. Ran, P. D. Hsu, J. Wright, V. Agarwala, D. A. Scott, F. Zhang, Genome engineering using the CRISPR-Cas9 system. Nature protocols 8, 2281–2308 (2013); published online EpubNov (10.1038/nprot.2013.143).

