## Supplemental data for "Regulation of *KRAS4A/B* splicing in cancer stem cells by the RBM39 splicing complex"

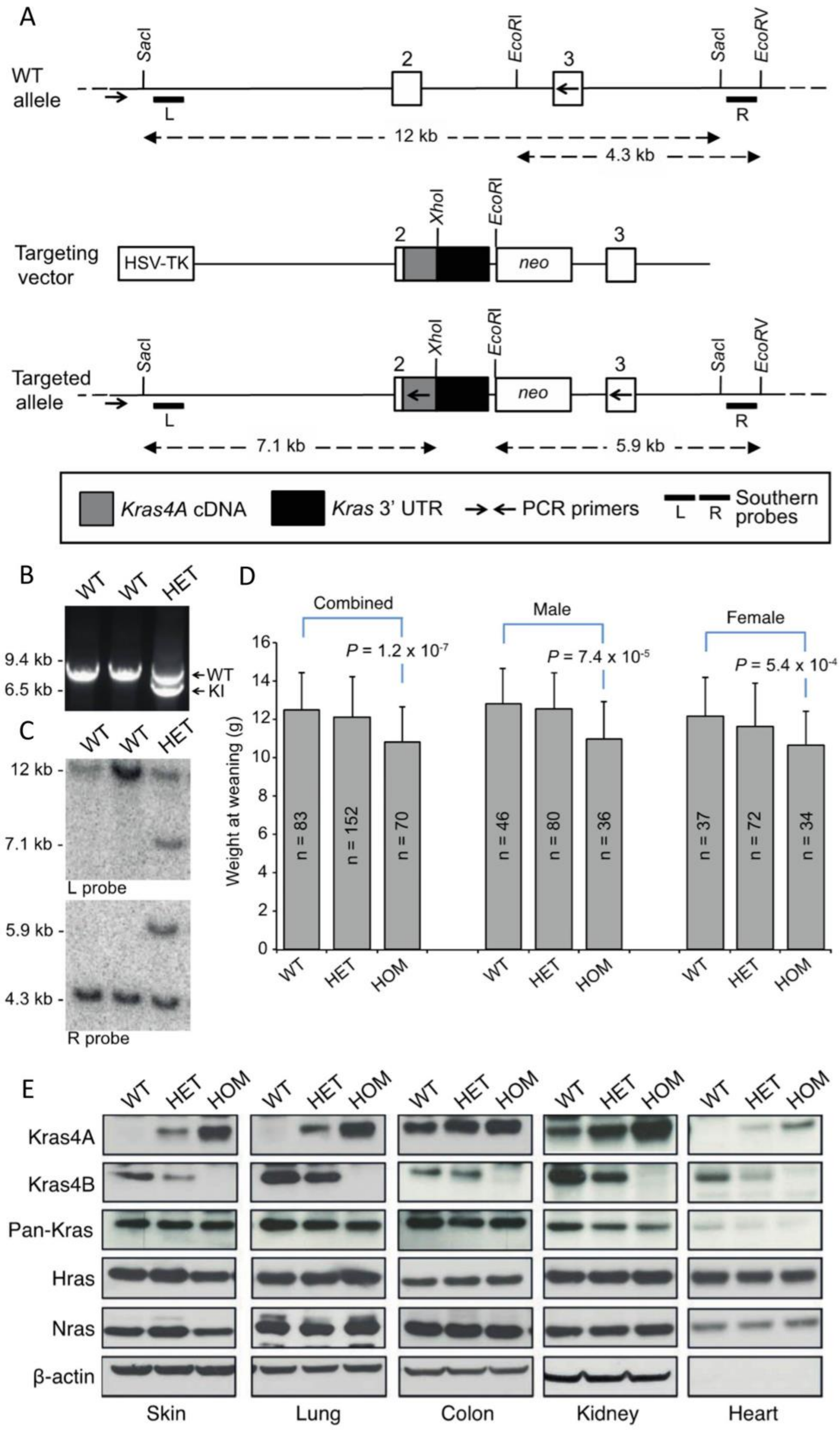

A
WT allele
SacI
2
EcoRI
3
SacI
EcoRV
L
R
12 kb
4.3 kb
Targeting vector
HSV-TK
2
XhoI
EcoRI
neo
3
Targeted allele
SacI
2
XhoI
EcoRI
neo
3
SacI
EcoRV
L
R
7.1 kb
5.9 kb
Kras4A cDNA
Kras 3' UTR
PCR primers
L R Southern probes
B
WT WT HET
9.4 kb
6.5 kb
WT
KI
C
WT WT HET
12 kb
7.1 kb
L probe
5.9 kb
4.3 kb
R probe
D
Combined
Male
Female
P = 1.2 x 10-7
P = 7.4 x 10-5
P = 5.4 x 10-4
Weight at weaning (g)
n = 83
n = 152
n = 70
n = 46
n = 80
n = 36
n = 37
n = 72
n = 34
WT HET HOM
E
WT HET HOM
Kras4A
Kras4B
Pan-Kras
Hras
Nras
β-actin
Skin
Lung
Colon
Kidney
Heart

**Figure S1**. Generation and characterization of mice that only express *Kras4A* from the *Kras* locus (*Kras4B-/-*). *(A)* Targeting strategy for the generation of a *Kras* allele with *Kras4A* cDNA knocked into the endogenous locus. *Kras4A* cDNA spanning a portion of exon 2 through exon 4A (in grey) and 3' UTR (in black) was homologously inserted into the *Kras* locus between exons 2 and 3. Locations of restriction endonuclease sites and fragment sizes, PCR screening primers and Southern probes are indicated. *(B-C)* PCR reactions *(B)* and Southern blots *(C)* show the expected DNA fragment sizes for successful targeting of the *Kras4A* allele. *(D)* Weights of WT, *Kras4B* heterozygous (HET) and homozygous *Kras4B-/-* (HOM) mice at weaning reveal a mild but significant reduction in weight of homozygous animals. This effect was not gender specific. *(E)* Western blot analysis of Kras4A, Kras4B, total Kras (Pan-Kras), Hras, and Nras across five normal tissues reveals expected changes in Kras4A and Kras4B between genotypes, and comparable levels of total Kras, Hras, and Nras across genotypes. β-actin is included as a loading control, although its expression is not detected in heart. Results were consistent across multiple mice from mixed B6/129 as well as pure FVB/N backgrounds. HET and HOM indicate *Kras4B* heterozygous and homozygous knockout genotypes, respectively.

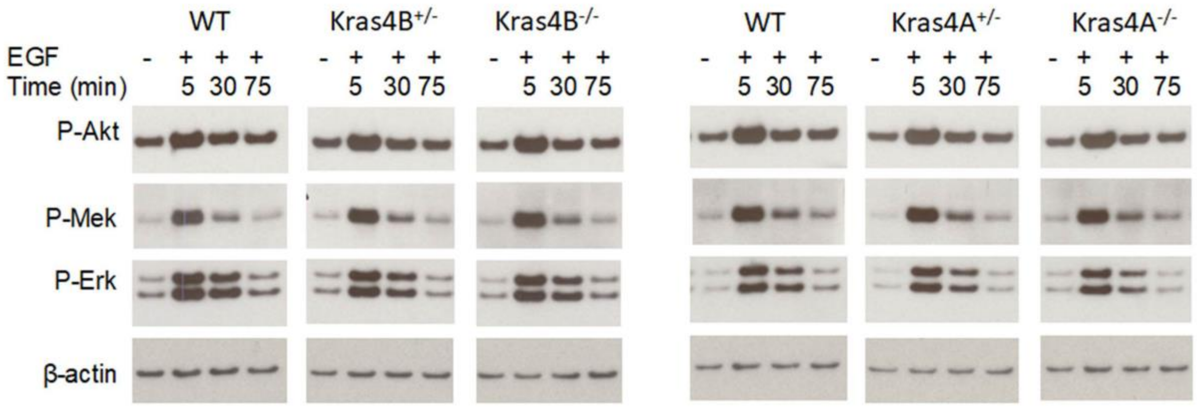


**Figure S2**. Effect of Kras isoforms on EGF-induced downstream signaling effectors. Analysis of phosphorylation of Akt, Mek and Erk in WT, Kras4A and Kras4B knockout MEFs showed elevated levels by EGF at the 5-minute time point. No apparent differences in EGF-induced activation of Akt, Mek and Erk were observed among mice of the indicated genotypes.

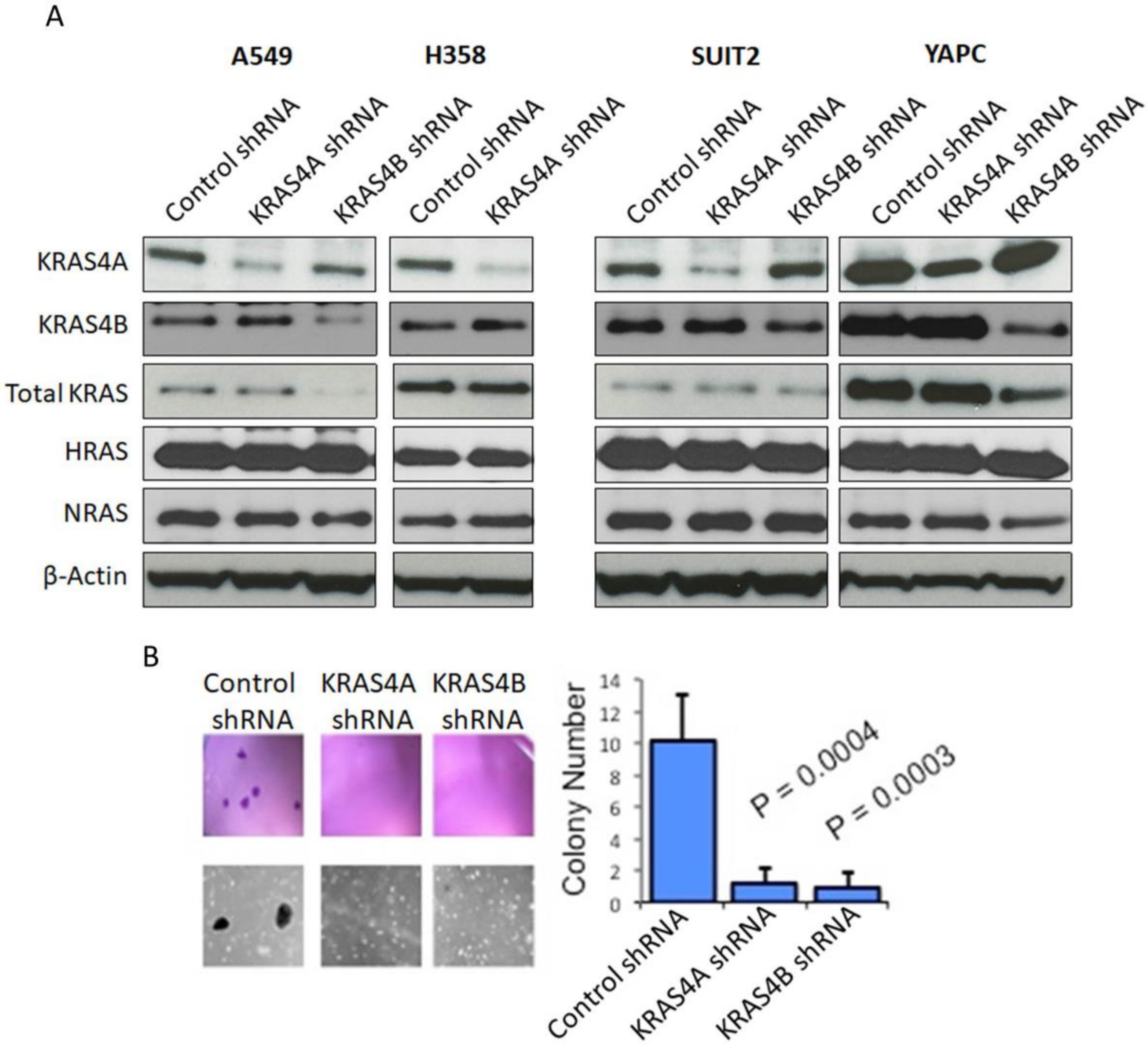


**Figure S3**. Ras protein levels in KRAS4A and KRAS4B knockdown human cancer cells. Lung cancer cells, A549 and H358, and pancreatic cancer cells, SUIT2 and YAPC, were transduced with indicated targeting shRNA, followed by puromycin selection. (A) RAS levels of the protein extracts from these selected cells with shRNA transfection were determined by western blotting. (B) The growth of SUIT2 cells with shRNA were determined by soft agar assay.

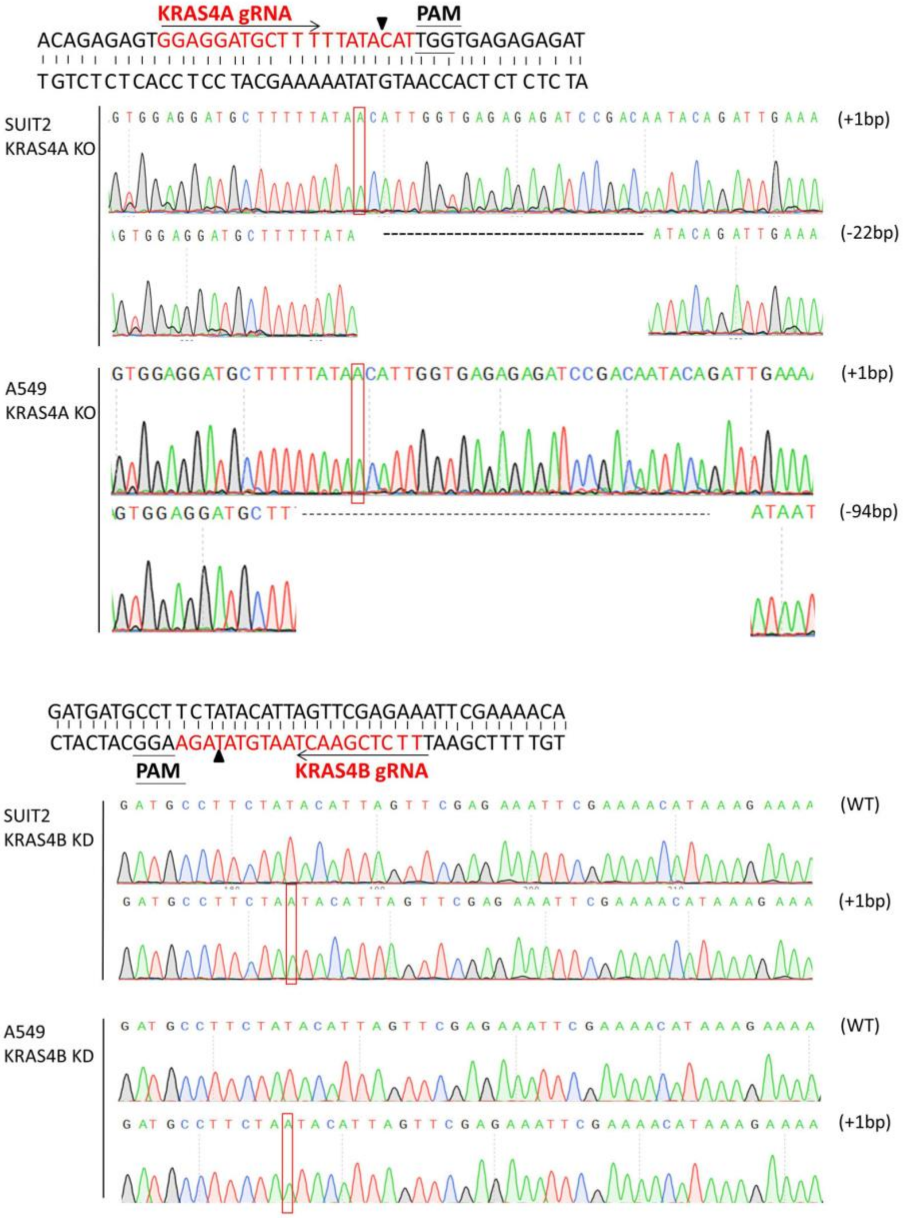


**Figure S4**. CRISPR/Cas9-based editing of the KRAS4A and KRAS4B exons. The actual indels of each allele in KRAS4A knockout and KRAS4B knockdown cells were determined by sanger sequencing.

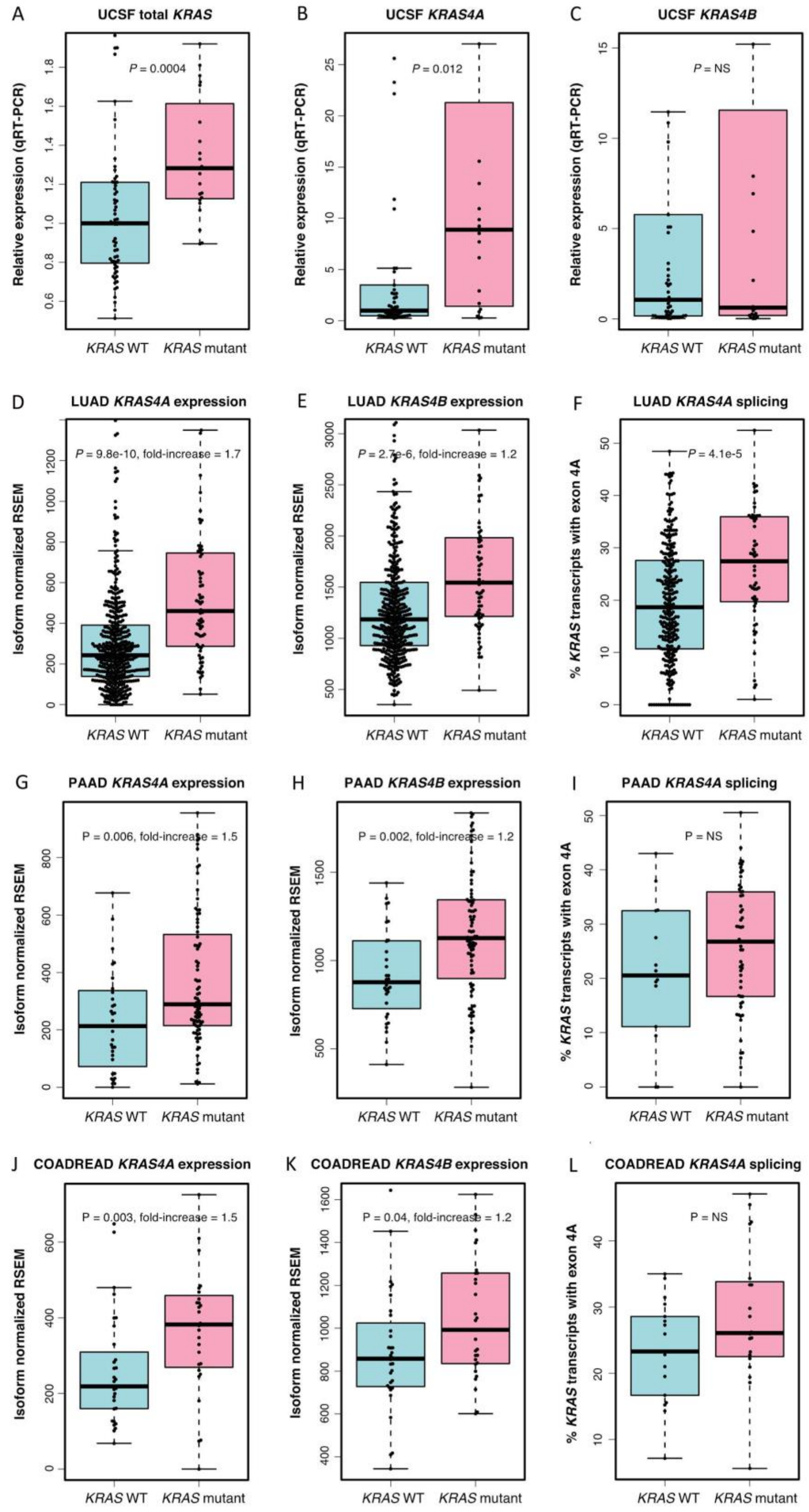


**Figure S5**. *KRAS4A* and *KRAS4B* expression in cancer. Total *KRAS* (A), *KRAS4A* (B), and *KRAS4B* (C) expression levels in human lung cancer were determined by TaqMan assay (n=63

for WT and n=23 for mutant KRAS ). The expression levels of *KARS4A* and *KRAS4B,* and the RNA seq reads of KRAS4A transcripts were assessed in lung adenocarcinoma (D-F), pancreatic adenocarcinoma (G-I) and colorectal adenocarcinoma (J-L) were assessed from TCGA datasets.

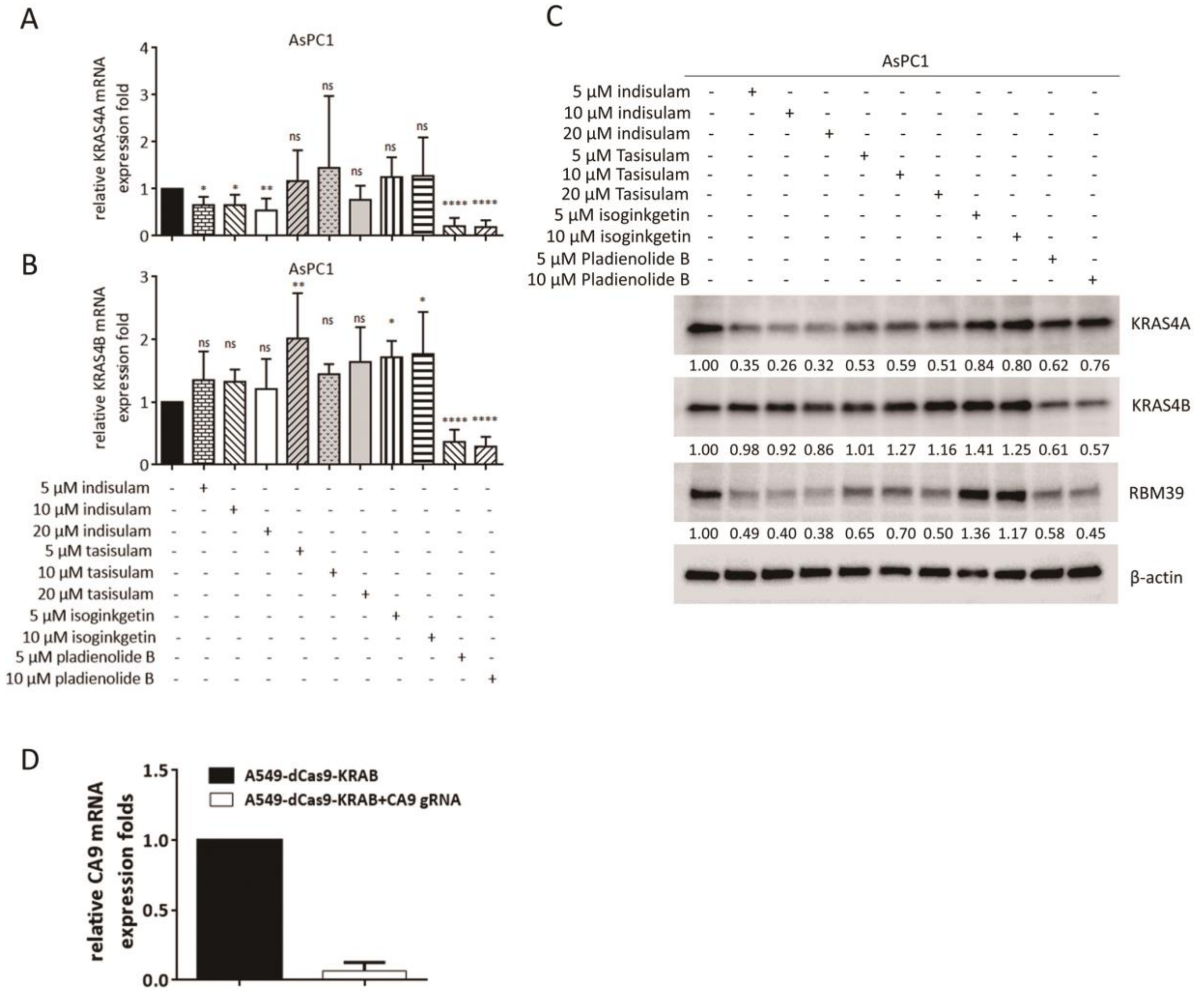


**Figure S6.** The RBM39 RNA-binding protein mediates *KRAS4A* splicing. The *KRAS4A* (A) and *KRAS4B* (B) mRNA levels were assessed by TaqMan analysis in AsPC1 cells after small molecule inhibitor treatment for 48hr. The specific inhibitors used and their concentrations are shown below the plots. Data are presented as mean ± s.e.m for four biological replicates. $*P < 0.05$; $**P < 0.01$ by one-way ANOVA with Tukey's multiple comparison. (C) The KRAS4A, KRAS4B and RBM39 protein levels were assessed in AsPC1 by Western blotting after small molecule inhibitor treatment for 48hr. Quantification of KRAS4A, KRAS4B and RBM39 levels was carried out using imageJ software. (D) Two sgRNAs targeting CA9 were transfected into BFP+A549 cells stably expressing dCas9-KRAB and then analyzed by Taqman analysis.

Supplementary Table 1: *Kras* mutations in carcinogen induced lung tumors from Kras4A heterozygous mice.

| | *Kras* Q61R | *Kras* Q61L | *Kras* Q61H | *Kras* G12D |
|---|---|---|---|---|
| 5 doses of urethane | 8/13 (61.5%) | 4/13 (30.8%) | - | - |
| 3 doses of urethane | 15/36 (41.7%) | 12/36 (33.3%) | 6/36 (16.7%) | 1/36 (2.8%) |
| 3 doses of MNU | 1/63 (1.6%) | - | 1/63 (1.6%) | 53/63 (84.1%) |

All tumors are from control *Kras4A* heterozygous mice as shown in Figure 1E. No tumors were found in carcinogen treated *Kras4A/Kras4B* double heterozygotes.
